# Two-brain microstates: A novel hyperscanning-EEG method for quantifying task-driven inter-brain asymmetry

**DOI:** 10.1101/2024.05.06.592342

**Authors:** Qianliang Li, Marius Zimmermann, Ivana Konvalinka

## Abstract

**Background:** The neural mechanisms underlying real-time social interaction remain poorly understood. While hyperscanning has emerged as a popular method to better understand inter-brain mechanisms, inter-brain methods remain underdeveloped, and primarily focused on inter-brain synchronization (IBS).

**New method:** We developed a novel approach employing two-brain EEG microstates, to investigate neural mechanisms during symmetric and asymmetric interactive tasks. Microstates are quasi-stable configurations of brain activity that have been proposed to represent basic building blocks for mental processing. Expanding the microstate methodology to dyads of interacting participants enables us to investigate quasi-stable moments of inter-brain synchronous and asymmetric activity.

**Results:** Conventional microstates fitted to individuals were not related to the different interactive conditions. However, two-brain microstates were modulated in the observer-actor condition, compared to all other conditions where participants had more symmetric task demands, and the same trend was observed for the follower-leader condition. This indicates differences in resting state default-mode network activity during interactions with asymmetric tasks.

**Comparison with existing methods:** Hyperscanning studies have primarily estimated IBS based on functional connectivity measures. However, localized connections are often hard to interpret on a larger scale when multiple connections across brains are found to be important. Two-brain microstates offer an alternative approach to evaluate neural activity from a large-scale global network perspective, by quantifying task-driven asymmetric neural states between interacting individuals.

**Conclusions:** We present a novel method using two-brain microstates, including open-source code, which expands the current hyperscanning-EEG methodology to measure and potentially identify both synchronous and asymmetric inter-brain states during real-time social interaction.

## 1 Introduction

To fully understand social cognition between individuals interacting together, it has been argued that it is inadequate to only measure neural processes in single, isolated individuals. Instead, a second-person approach should be employed to capture both the neural processes within individuals engaged in interaction, but also the dynamic interactions and processes between individuals (Schilbach et al., 2013; Konvalinka and Roepstorff, 2012; Dumas, 2011; De Jaegher et al., 2010; Dingemanse et al., 2023). The interpersonal and inter-brain mechanisms tap into important aspects of social processes that evolve over time between interacting partners, as well as more fine-grained mechanisms of coordination (Konvalinka and Roepstorff, 2012; Konvalinka et al., 2023). In the last two decades, the simultaneous measurement of brain activity from multiple individuals (using multi-person EEG, MEG, fNIRS or fMRI recordings) has become one particular trend that has emerged in quantifying the interpersonal, dynamic neural mechanisms between interacting individuals, and has been coined hyperscanning (Montague et al., 2002). A growing body of hyperscanning studies has shown that people’s brain rhythms or activities synchronize when they engage in social interaction with each other (see Babiloni and Astolfi (2014); Czeszumski et al. (2020) for reviews), while e.g., coordinating or synchronizing their actions (Dumas et al., 2010; Zamm et al., 2018; Lindenberger et al., 2009), cooperating with one another (Hu et al., 2018; Cui et al., 2012), turn-taking in verbal interaction (Ahn et al., 2018), or while merely able to see each other in unstructured tasks (Koul et al., 2023). However, while hyperscanning has been around for two decades, with growing efforts to standardise the methodology (Ayrolles et al., 2021), the inter-brain analysis methods remain underdeveloped, lacking clear standards (Zamm et al., 2024; Zimmermann et al., 2024), and primarily focused on measuring synchrony between people’s neural signals, drawing critique with respect to the underlying functional significance of inter-brain synchrony (IBS) (Hamilton, 2021; Holroyd, 2022). Hence, more innovative inter-brain methods are required to go beyond IBS, to better understand functional neural mechanisms that underlie different types of social processes and interactive outcomes, while still taking advantage of simultaneous neural recordings in interacting individuals.

Here, we employ a new approach to investigate two-brain mechanisms, deviating from the common approach of measuring functional connectivity between brains (Xu et al., 2024) which assumes that two brains in interaction synchronize and have functionally meaningful connectivity patterns that may even causally facilitate social interactions (Novembre and Iannetti, 2021). The main motivation is to be able to identify spatiotemporal interpersonal asymmetries in neural activity patterns, which may emerge in real-time social interactions but are still related to functionally relevant neural processing in individual brains i.e., processes that enable or underlie different social tasks, roles, and interaction strategies between people when they interact with each other. Asymmetric interaction behaviour may include e.g., taking on leader or follower roles when coordinating actions, producing complementary rather than imitative actions, initiating versus mimicking behaviour, or observing versus being observed (Sebanz et al., 2006; Konvalinka et al., 2023). Inter-brain synchrony may still underlie such asymmetric behaviour, or may exhibit asymmetric functional connectivity patterns, e.g., directed coherence between different brain regions of the two interacting partners (Babiloni et al., 2007; Astolfi et al., 2010). However, there may be important functionally relevant differences in neural processes between participants with different roles in e.g., action-perception coupling, which are still poorly understood, and would help to better elucidate the neural basis of real-time social interaction.

For example, previous work has shown that asymmetric leader-follower roles emerge when people interact with one another, and that they can be predicted based on 10 Hz oscillations measured using dual-EEG, when employing two-brain analyses (Konvalinka et al., 2014). In this study, participants engaged in an interactive finger-tapping task with each other or with a computer metronome. When applying multivariate decoding techniques to the two-brain data, the 10 Hz power across only the leaders’ frontal electrodes were picked up as reliable classifiers of interactive versus non-interactive conditions. As leaders had to suppress the sounds they heard from the other person and focus more on monitoring their own taps, this suggests that suppression of frontal 10 Hz power may be a potential neural mechanism underlying leading, or increased self-monitoring, behaviour. Meanwhile, followers exhibited more similar frequency power modulations between interactive and non-interactive conditions, given that they were following the other person or the computer, respectively. Notably, this neural asymmetry was only picked up when using dual-EEG data, rather than focusing on individual neural modulations.

Another study investigating role-differences between senders and receivers in a cooperative game found an asymmetry in alpha and low beta event-related desynchronization between the senders and receivers (Flösch et al., 2023). Here, the participants engaged in a cooperative Pacman Game, where one participant (the sender) sent information about the correct moving direction in the form of a picture cue to the other player (the receiver), who had to correctly decode the picture cue in order to move towards the goal. The receiver exhibited a larger alpha/beta power decrease compared to the sender, potentially indicating higher cognitive demands, in line with the role asymmetry which required the receiver to represent more information than the sender.

In addition to having different roles in social interaction, such as that of a leader and follower, or sender and receiver, people may also adopt different interpersonal strategies. For example, people may choose to adapt to each other as mutual followers, or mutually decide to ignore one another, exhibiting so-called leader-leader behaviour (Heggli et al., 2021). Such different interpersonal strategies during real-time social coordination have been shown to emerge between musicians, and have different underlying neural networks. E.g., mutual adaptation exhibits more frequent occurrence of phase-locked activity in the alpha frequency band within the right temporoparietal network compared to leader-leader behaviour, presumably showing more recruitment of action-perception neural networks. However, these differences were found using single-brain analyses; hence, it is unclear whether there were additional inter-brain asymmetries.

In this paper, we were interested in exploring both synchronous as well as asymmetric neural spatiotemporal mechanisms during social interaction, by developing a novel two-brain microstates method, which could be freely applied to interaction conditions that involve both symmetric and asymmetric tasks and roles. Microstates are quasi-stable configurations of brain activity that have been reliably replicated across studies, and proposed to be basic buildings blocks for mental processing (Lehmann et al., 1987; Michel and Koenig, 2018). Each microstate topographical map is thought to be generated by the coordinated activity of multiple neural sources, thus transitions between microstates may be interpreted as switching between different neural networks of the brain (Khanna et al., 2015; Michel and Koenig, 2018). These underlying neural networks can thus be investigated by quantifying the spatiotemporal dynamics of the microstates, e.g., by looking at the average lifespan of a microstate (duration), the relative tendency for a specific microstate to be active relative to the others (coverage), or the transition probabilities between the microstates. More complex temporal dynamics can also be estimated, such as entropy, Lempel-Ziv complexity and Hurst exponents (von Wegner et al., 2023). These microstate dynamical features can in turn be correlated to various functional roles, and a growing number of studies have employed microstates to investigate the neural mechanisms of cognitive processes and clinical disorders, e.g. schizophrenia, psychosis, Alzheimer’s disease, epilepsy, bipolar disorder, depression, autism spectrum disorder, and many others (reviewed in Khanna et al. (2015); Michel and Koenig (2018); Tarailis et al. (2023); Schiller et al. (2024)).

Traditionally, microstates have been primarily applied to resting-state data, but a minority of studies have also applied microstates to task-related EEG (see Schiller et al. (2024) for a review of microstates applied to social or affective tasks) to e.g., decode emotions during video watching (Liu et al., 2023), pinpoint the articulatory onset based on event-related potentials (Jouen et al., 2021), and as a biomarker for stable motor output during a force production task (Pierpaolo et al., 2022). In this study, we expanded the microstate methodology to dyads of interacting participants (two-brain microstates), which enables us to investigate quasi-stable moments of inter-brain synchronous activity, while not constraining the quantification of inter-brain synchrony to symmetric brain states. In other words, a single two-brain microstate could be asymmetric for the two participants in a dyad, if that particular combination of asymmetric brain states often occurred simultaneously in the dyads. This feature enables two-brain microstates to potentially uncover contrasting patterns of brain activity when people engage in symmetric or asymmetric interactive tasks.

To evaluate the functional role of two-brain microstates, we applied this new method to hyperscanning EEG data from an experiment employing the adapted mirror game paradigm (Zimmermann et al., 2022). The mirror game paradigm engages pairs of participants in a joint movement game, where they are asked to improvise motion together by creating synchronized and interesting movements (Noy et al., 2011). The participants had to perform horizontal finger movements by moving a slider, and we manipulated whether the participants could see each other’s hand. The participants thus produced horizontal movements across a number of conditions involving symmetric, but not necessarily interactive tasks: independent movements (uncoupled condition), interactive movements (coupled condition), similar but not interactive movements while both synchronized to the same pre-recorded signal (control condition). They also engaged in two asymmetric interactive conditions where they either had asymmetric roles or asymmetric tasks: where one was assigned the role of a leader and the other of a follower (follower-leader condition), and where one was told to improvise independent movements while observed by the other (observer-actor condition). Figure 1 shows a schematic of the experiment.

**Fig. 1.**
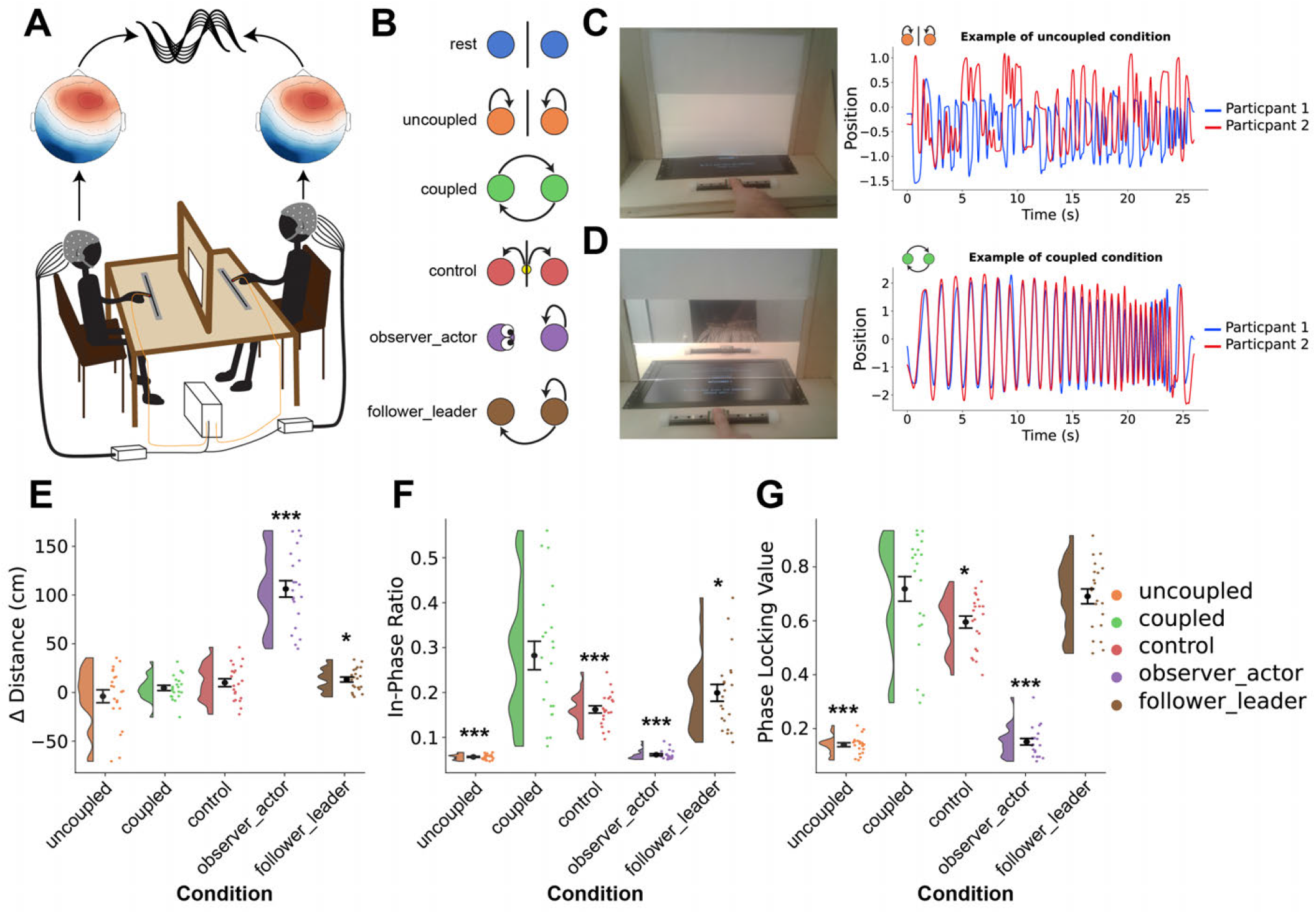
Overview of the mirror-game setup and the different behavioural conditions. A) Dyads performed the mirror-game paradigm, during which we measured their EEG. The two-brain EEG data were subsequently analyzed using microstate analysis and inter-brain synchronization was estimated. B) The dyads performed four types of symmetric (rest, uncoupled, coupled and control) and two asymmetric (observer-actor and follower-leader) conditions. C) Example of the non-interactive uncoupled condition and the finger positions of each participant. D) Example of the interactive coupled condition, with strong behavioural synchronization. E-G) Behavioural performance measures for each condition. All statistical tests were performed in contrast to the principal interactive coupled condition. E) The asymmetric conditions had a greater difference in distance moved compared to the coupled condition, indicating the actor and leader moved more than the observer and follower, respectively. The coupled condition displayed the highest behavioural synchrony, as measured by F) the ratio of time the finger movements were in-phase (within 10^*°*^ from each other) and G) the phase locking value, followed by the follower-leader and control conditions, with uncoupled and observer-actor conditions displaying minimal synchrony. Adapted from Zimmermann et al. (2022).

Previously, we conducted intra-brain analyses on this data set, and reported unique individual behavioural and neural signatures when performing actions when observed by others (Zimmermann et al., 2022). Zimmermann et al. (2022) showed that participants produced movements that were slower, less variable, but more exaggerated in amplitude, when they were observed by the other person, compared to when they were unable to see each other, indicating audience effects. In addition, observed actions were characterized by increased widespread functional connectivity in the alpha frequency range in contrast to both individual (uncoupled) and interactive (coupled) actions, potentially indicating increased self-monitoring and focus.

In order to explore inter-brain mechanisms across the different symmetric and asymmetric interaction conditions, we applied our novel two-brain EEG microstates method. We hypothesized that the coupled condition would lead to higher inter-brain synchrony than the other conditions (particularly the non-interactive ones), based on previous research; and that asymmetric conditions (follower-leader, observer-actor) would lead to more pronounced asymmetries between the two persons’ spatiotemporal neural dynamics compared to the coupled condition, which would be captured with two-brain microstate dynamics. Crucially, we compared whether microstate dynamics would be better differentiated across the different interaction conditions when they were fitted to two-brains rather than individual brains.

## 2 Materials and Methods

### 2.1 Microstate analysis

Microstates are typically estimated in broadband, but a recent paper (Férat et al., 2022) demonstrated the value of spectrally specific microstates. In addition, given previous literature showing that alpha and beta oscillations are modulated during motor interactions (Conway et al., 1995; Perez et al., 2006; Klimesch et al., 2007; Tognoli et al., 2007; Baker, 2007; Dumas et al., 2012), we estimated the microstates in the alpha (81-3 Hz), beta (13-30 Hz) and broadband (1-40 Hz) frequency ranges. Single-brain microstate analysis was performed using the open-source EEG microstate python package by von Wegner and Laufs (2018). Briefly, EEG from all individuals were concatenated along the time axis, in order to fit the microstates over all the subjects and obtain the same topographic maps. Global field power (GFP) was computed and EEG topographies at local GFP maxima were clustered using the modified *K*-means algorithm (Pascual-Marqui et al., 1995). We ran *K*-means 100 times with 3 to 10 clusters and used the cross-validation criterion (Pascual-Marqui et al., 1995) to determine the final number of clusters. The determined microstate topographies were competitively fitted back by assigning each EEG data time point one microstate label based on maximal correlation, to yield time series of microstate sequences. The following commonly used microstate features were estimated: ratio of time covered (relative time a microstate is active as a ratio of the total time), duration (mean duration a given microstate remains stable, i.e. occurs consecutively), occurrence (mean number of times a microstate occurred during a one second period), transition matrices (mean probability of transitioning from one microstate to itself or another) and entropy (Shannon entropy) were computed for the microstates (von Wegner and Laufs, 2018; Tarailis et al., 2023). Figure 2 shows a flowchart of the microstate methodology.

**Fig. 2.**
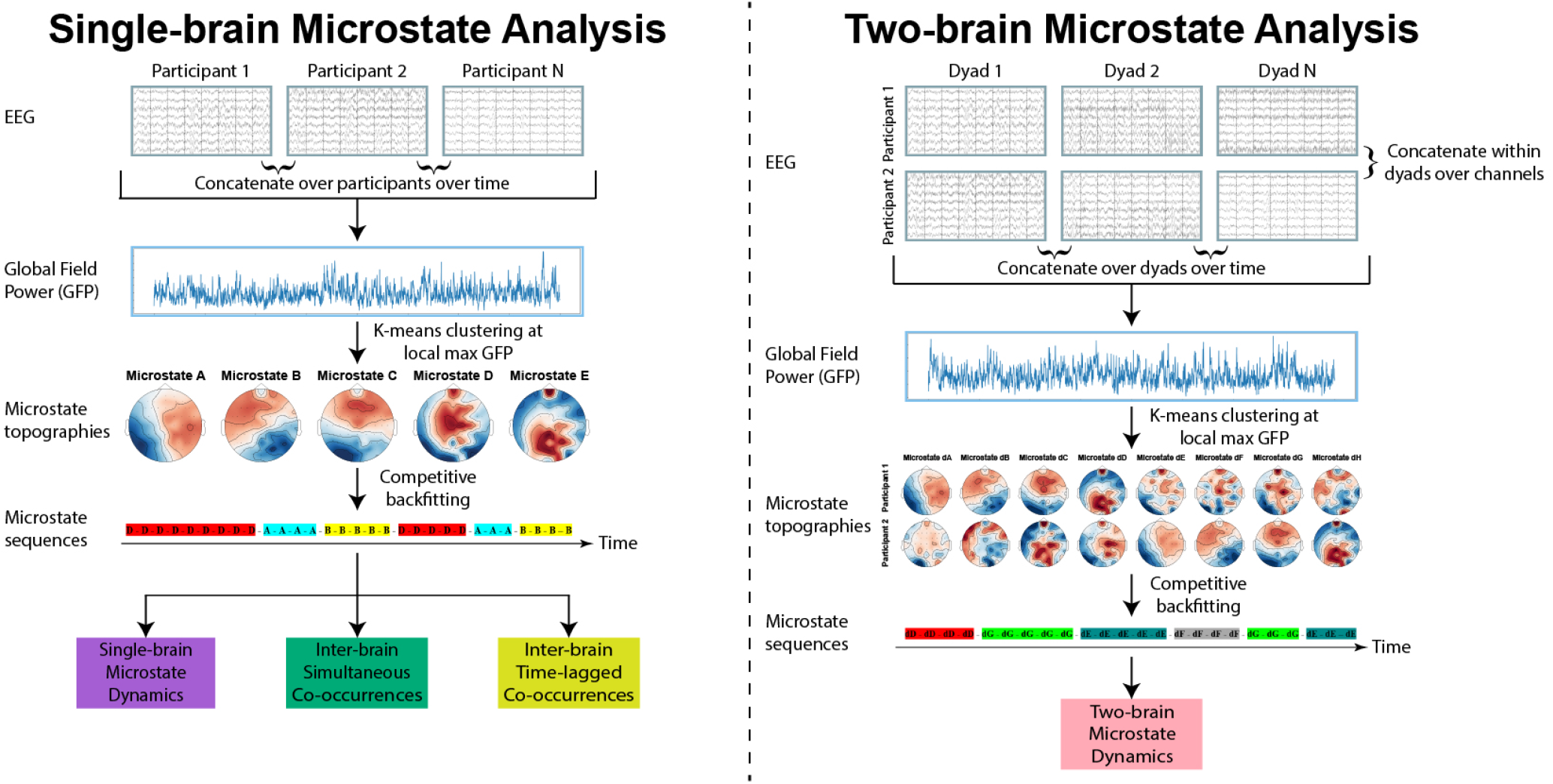
Flowchart of the microstate analyses. The left column shows the classical single/intra-brain microstate analysis. In addition to the single-brain microstate dynamics, we also estimated the dynamics of co-occurring inter-brain microstates based on single-brain microstates fitted to each individual. The right column shows the two-brain microstate analysis.

One definition of inter-brain synchrony is that the two brains are displaying similar activity at the same time. To investigate whether the participants of the mirror-game exhibited this form of synchronous activity, we quantified how often the dyads were in the exact same single-brain microstates at the same time, and looked at the dynamics of this co-occurrence of quasi stable global brain activity for the different behavioural conditions. Specifically, we created an inter-brain microstate sequence time series based on the intersection at each time point of the two individual microstate sequence time series from a dyad. In other words, if both brains are in microstate A at the same time point, then the inter-brain microstate sequence time series at that time point will also be A, however if their labels are different, then they do not intersect and we labelled it arbitrarily as Z to denote zero inter-brain synchrony.

However, the criteria of being in the exact same global neural activity state might be too strict, so we also worked with two less constrained definitions of inter-brain synchrony. In the first implementation, we allowed for time-lags to shift either participant 1 or 2’s microstate sequence time series with up to 1 second in both directions, in order to determine a specific lag that had the highest amount of time-shifted interbrain synchrony (comparable to cross-correlation).

In the second implementation we removed the constraint of the two brains in a dyad being in the exact same state, and as long as the two potentially different states consistently cooccurred over time, we considered them to be synchronous. This definition of inter-brain synchrony gave rise to what we refer to as two-brain microstates, where each two-brain microstate consists of a fixed global neural activity pattern, which can be different for the two individuals. To implement it in practice, we concatenated the EEG data from each pair along the channel axis, before concatenating all the dyads along the time axis. By concatenating over the channel axis as opposed to the time axis for the two participants in a dyad, we allowed the two-brain microstates to have different topographies for each person in a dyad as long as they occurred simultaneously, i.e., a two-brain microstate could be asymmetric. Note that a two-brain microstate thus consists of activity in 128 channels, but for visualization we plotted the electrodes corresponding to each individual separately to show their topographies, but it is in fact one “inter-brain state”. Additionally, there was no issue of electrode adjacency when concatenating over the channel axis, as the *K*-means algorithm does not take into account the order of the dimensions (channels) of the data. To retain the polarity invariance in two-brain microstates, we computed the spatial correlation for all four combinations of possible polarity configurations for the two brains, and used the highest correlation (see discussion for more details). The above-mentioned common single-brain microstate features were also estimated for two-brain microstates, to investigate joint spatiotemporal neural dynamics. In addition, we also estimated long-range temporal correlations using detrended fluctuation analysis (DFA) to compute the Hurst exponent (Hardstone et al., 2012) (see supplementary materials for more information). Figure 2 shows a flowchart of the two-brain microstate methodology in comparison to the single-brain microstate methodology.

### 2.2 Source localization

We utilized eLORETA, as implemented by MNE-Python v1.3.1 (Gramfort et al., 2013), to obtain cortical current estimates for the two-brain microstates. The FreeSurfer average brain template from FreeSurfer 6 (Fischl, 2012) was used to construct the boundary element head model and forward operator for the source modelling. The regularization parameter was set to *λ*^2^ = 1*/*9 and we used the normal orientation, i.e. only the portion of the activity perpendicular to the cortex. The time series of the 20484 source vertices were further collapsed into 68 cortical patches based on the Desikan Killiany atlas, by first aligning the dipole orientations by shifting vertices with opposite polarity to the majority of vertices by *π*, followed by averaging the amplitudes of all vertices within a patch. The phase shifting prevents the vertices with opposite polarities from canceling each other out during the averaging operation.

### 2.3 Mirror-game dataset

The data we applied the two-brain microstate method to have been previously described in Zimmermann et al. (2022). Originally, 25 pairs were recruited to participate in a mirror-game paradigm while simultaneous EEG was recorded. The pairs consisted of 11 mixed-sex, and 14 same-sex pairs (1 female-female, 13 male-male). 16 pairs were friends/partners, and 9 pairs did not know each other before the experiment, but were introduced to each other during preparations of the experiment. Due to technical problems during the experiment and EEG data quality issues, four pairs were dropped, hence 21 pairs (42 participants; 26% female; aged 20-33, 24.36 years on average) were included in the analysis.

The mirror-game is an experimental paradigm designed for examining the dynamics of two interacting individuals (Noy et al., 2011), where the two participants improvise motion either alone or together with a partner. The participants were divided by a liquid crystal screen, which could be turned off during non-interactive conditions (Figure 1C) or on during interactive conditions (Figure 1D), manipulating whether participants had visual feedback of each other’s movements or not. The vision of their partner’s face and body was always blocked. The participants thus produced horizontal movements across a number of conditions involving symmetric, but not necessarily interactive tasks: 1) 2 min *rest* condition at the start and end of experiment (screen off), 2) 16 × 25 sec *uncoupled* movements where participants could not see each other and produced independent movements (screen off), 3) 16 × 25 sec *coupled* movements where participants could see each other and synchronized their movements (screen on), 4) 16 × 25 sec *control* condition where participants had to follow a dot on the screen (screen off), hence interacting with the same stimulus but not with each other (the movement of the dot corresponded to movements produced in the coupled condition of pilot participants), 5) 2 × 8 × 25 sec *observer-actor* with one participant producing improvised movements while observed by the (non-moving) partner (screen on), and 6) 2 × 8 × 25 sec *follower-leader* where each participant was designated a specific role, while able to see each other (screen on). Figure 1 shows a schematic of the experiment and more details can be found in the supplementary material and Zimmermann et al. (2022).

### 2.4 Movement data analysis

The finger movements were recorded using Polhemus LIBERTY and pre-processed as described in Zimmermann et al. (2022). To estimate the behavioural performance of each dyad, i.e., their movement synchronization, the following behavioural features were computed from their finger positions over time: the difference in distance moved between the interacting participants, the amount of time their movements were in-phase (defined as being within 10°), and the phase locking value (Lachaux et al., 1999). The Hilbert transform was utilized to estimate the phase of each person’s movement time series.

### 2.5 EEG preprocessing

The EEG was recorded and pre-processed as described in Zimmermann et al. (2022). Briefly, two synchronized 64channel Biosemi EEG set-ups were recorded at a 2 kHz sampling frequency, followed by bandpass filtering at 1-40 Hz, and downsampling to 256 Hz. Manual visual inspection was performed to clean the data, and independent component analysis (ICA) was used to correct for eye movements and eye blinks.

### 2.6 Statistical analysis

Two-tailed non-parametric permutation tests were used to investigate differences in group means between the principal interactive *coupled* condition with all other behavioural conditions. In order to discern whether any differences were specific for the interaction itself, we also created surrogate data in the form of pseudo-pairs and compared with the real-pairs. A pseudo-pair is an artificial pair created by pairing the data from one participant with partners from all the other pairs except the real partner. Thus, we created 21 × (21 *−* 1) = 420 pseudo-pair surrogate data, which consisted of data from artificial pairs with individuals that both performed a given behavioural task; however, they were not time-locked to each other due to the lack of interaction (e.g., the observer from one dyad was paired with the actor from another dyad). Multiple testing correction was performed using false discovery rate (FDR) and applied to each frequency range and feature type separately. The significance level was 0.05 for all hypothesis tests and the number of asterisks corresponded to different p-values (^*^*p* < 0.05, ^**^*p* < 0.01, ^***^*p* < 0.001). Results are shown as mean with standard error of the mean (SEM).

## 3 Results

### 3.1 Interactive conditions were associated with greater behavioural synchrony

To evaluate the behavioural performances in the different conditions, we computed three movement features for each dyad and compared each condition with the principal interactive condition: the *coupled* condition. The asymmetric *observer-actor* (p-value < 0.001) and *follower-leader* (pvalue = 0.047) conditions exhibited a significantly greater difference in distance moved between the interacting participants compared to *coupled* condition, based on two-sided permutation tests with FDR multiple-comparison correction (Figure 1E). This difference indicates that the actor and leader moved their fingers more than the observer and follower, respectively. The interactive *coupled* condition displayed the greatest behavioural synchrony, evident by the high ratio of time the fingers of the dyads were in the same phase (Figure 1F) and high phase locking value (Figure 1G). The interactive *follower-leader* condition closely followed the *coupled* condition, with slightly lower in-phase ratio (Figure 1F; FDR corrected p-value = 0.029) and similar phase locking values (Figure 1G; FDR corrected p-value = 0.62). The nonsocial *control* condition was also associated with behavioural synchrony between the participants, due to the participants synchronizing to the same stimuli, albeit with a significantly lower in-phase ratio (Figure 1F; FDR corrected p-value < 0.001) and phase locking value (Figure 1G; FDR corrected p-value = 0.033) compared to the *coupled* condition. Lastly, the *uncoupled* and *observer-actor* conditions displayed minimal behavioural synchrony, as expected (Figure 1F-G; FDR corrected p-values < 0.001).

### 3.2 Single-brain microstate dynamics were not related to the mirror-game conditions

Applying the microstate methodology to all the 42 individual single-brain EEG data, filtered in the alpha frequency range, yielded five microstates explaining around 56% of the variance (Figure 3A). The corresponding microstates determined in the beta and broadband frequency can be found in Supplementary Figure S1A and Supplementary Figure S2A, respectively. The topographies of the determined microstates estimated in the alpha, beta and broadband frequency range were very similar with each other (Supplementary Figure S3), and also qualitatively similar to the conventionally found resting-state EEG microstates in the literature (Michel and Koenig, 2018; Tarailis et al., 2023; Koenig et al., 2023). Thus we sorted and labelled the microstates in line with the prototypes from literature with microstate A showing a leftright orientation, B with a right-left orientation, C with an anterior-posterior orientation, D with a fronto-central maximum, and E with an occipito-central maximum (Férat et al., 2022; Tarailis et al., 2023; Koenig et al., 2023).

**Fig. 3.**
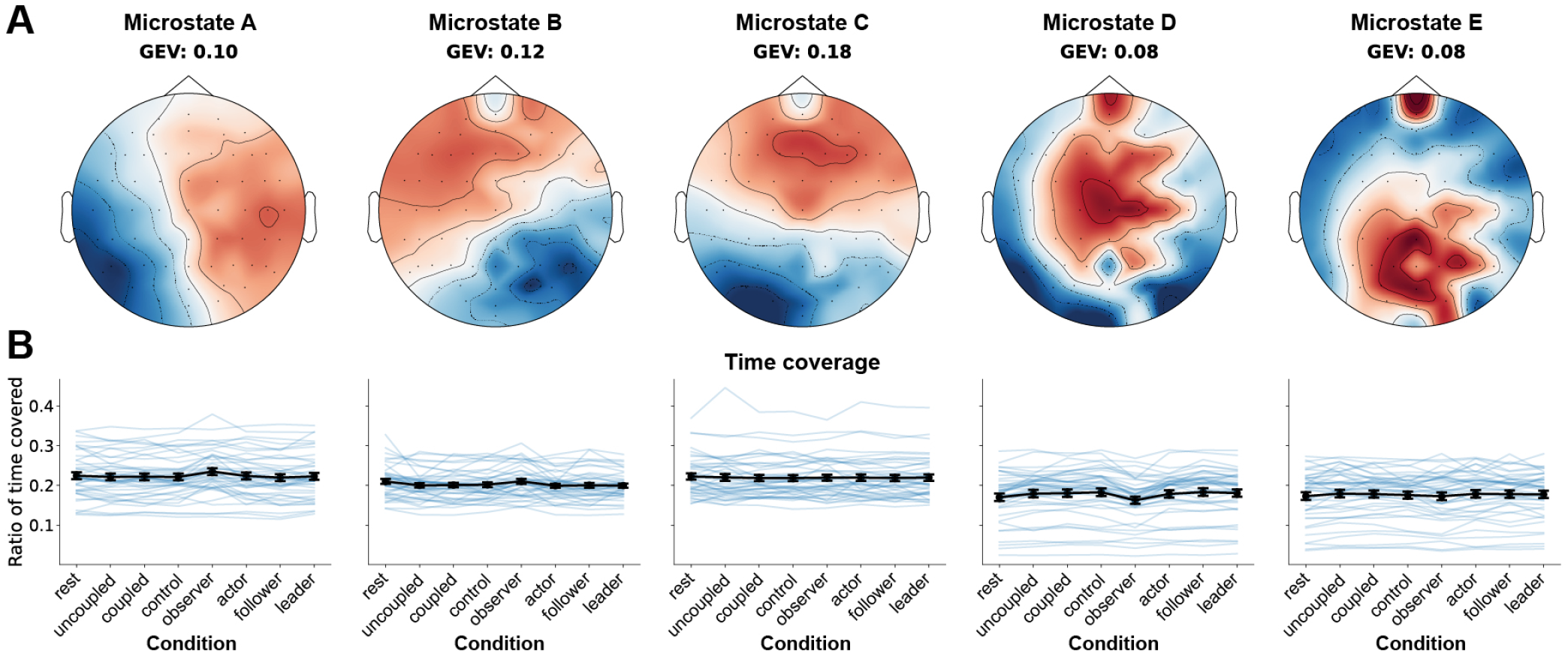
Conventional single-brain EEG microstate analysis. A) Microstate analysis was performed on the 42 individual EEG timeseries filtered in the alpha frequency range, and five microstates were determined, which explained around 56% of the variance. B) The ratio of time covered by each microstate was not related to the different behavioural conditions. GEV: global explained variance.

None of the computed alpha microstate features were related to the different behavioural conditions in the mirror-game, e.g., the ratio of time coverage of each of the five microstates was similar for all the different conditions (Figure 3B; lowest uncorrected p-value = 0.16, lowest FDR multiple test corrected p-value = 0.99, two-sided permutation test) and a similar result was also observed for the other microstate features: duration, occurrence, transition probabilities and entropy (Supplementary Figure S4, S5 and S8A), indicating no significant differences between any of the alpha microstate features across the conditions (lowest uncorrected p-value = 0.14, lowest FDR corrected p-value = 0.99). The same results were observed for the beta (Supplementary Figure S1, S6 and S8B; lowest uncorrected p-value = 0.015, lowest FDR corrected p-value = 0.76), and broadband frequency range (Supplementary Figure S2, S7 and S8C; lowest uncorrected p-value = 0.036, lowest FDR corrected p-value = 0.72) determined microstate features.

A high degree of correlations were found between the microstate features: duration, occurrence, transition probabilities and time coverage. For instance, duration of microstate A was highly correlated with occurrence, transition probability and time coverage of microstate A, and the same was observed for the other microstates (Supplementary Figure S9). For brevity, we focus the visualizations on time coverage (the other features can be found in the supplementary material).

To summarize, the single-brain microstates estimated from the mirror-game were qualitatively similar to the canonical resting-state microstates previously described. However, the dynamics of the microstates were not related to the mirrorgame conditions.

### 3.3 Inter-brain co-occurrences of microstates were not related to the mirror-game conditions

Estimation of inter-brain synchrony, defined as the cooccurrence of the same microstate at the same time point, also revealed no differences associated with the mirror-game conditions for time coverage of alpha microstates (Figure 4; lowest uncorrected p-value = 0.20, lowest FDR multiple test corrected p-value = 0.99, two-sided permutation tests). A similar result was also observed for the other alpha microstate features: duration, occurrence, transition probabilities and entropy with no significant associations between the inter-brain microstate features and the mirror-game conditions (Supplementary Figure S10, S11 and S12A; lowest uncorrected p-value = 0.11, lowest FDR corrected p-value = 0.98). A lack of association between inter-brain microstate dynamics and mirror-game conditions was also observed for beta (Supplementary Figure S13, S14 and S12B; lowest uncorrected p-value = 0.02, lowest FDR corrected p-value = 0.86) and broadband microstate dynamics (Supplementary Figure S15, S16 and S12C; lowest uncorrected p-value = 0.06, lowest FDR corrected p-value = 0.98).

**Fig. 4.**
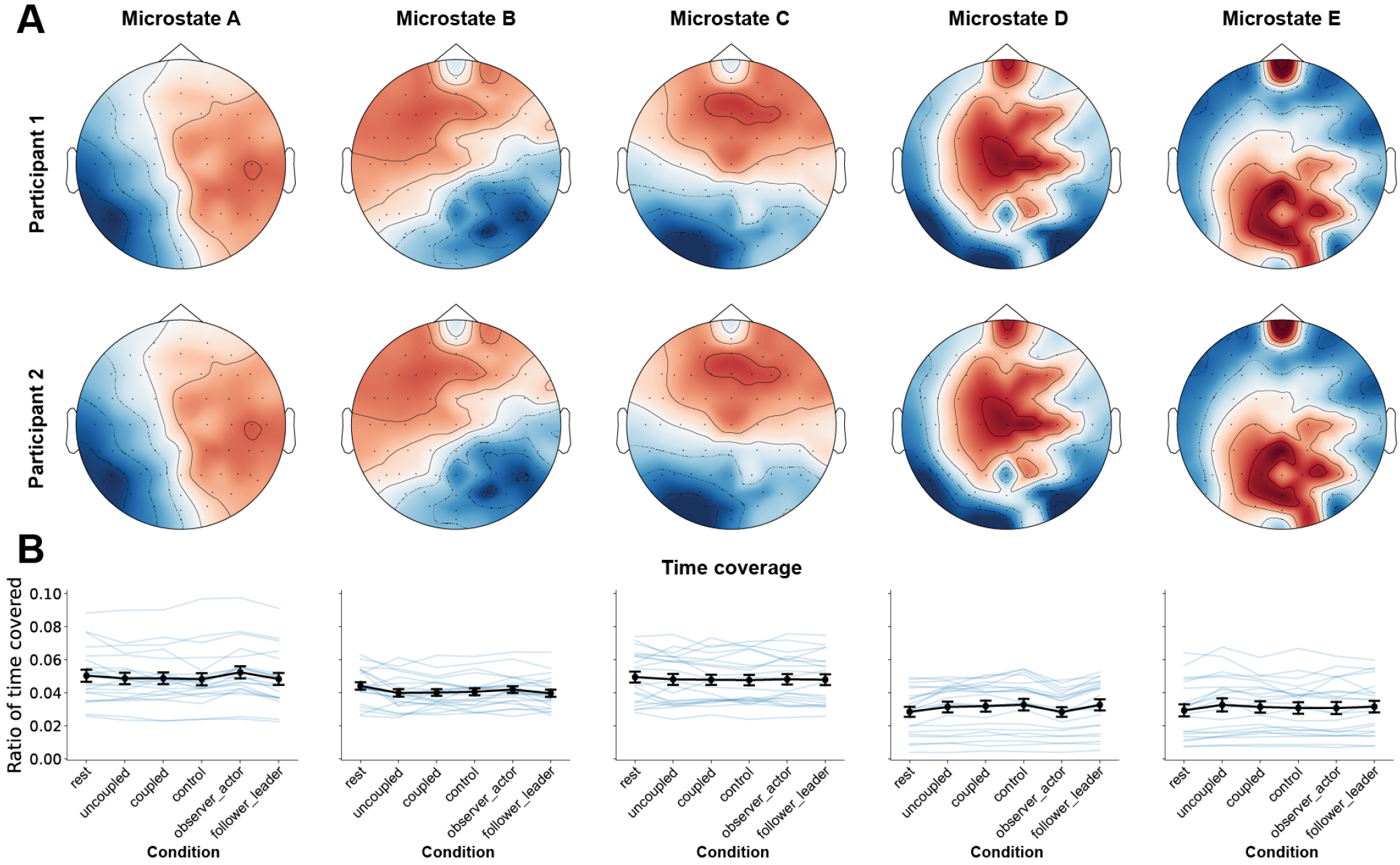
Inter-brain synchrony of microstates. A) We investigated the co-occurrence of similar microstates in the two participants in the dyads performing the mirror-game. B) The ratio of time the participants were in the same microstate was not related to the different behavioural conditions. The ratio of time coverage was normalized to the total time they were in the same microstate and the time not in the same microstate (Supplementary Figure S17), hence the low values.

Surprisingly, the amount of time the two participants in a dyad were in the same microstate was not significant from chance-level for alpha, beta, and broadband microstates (Supplementary Figure S17; lowest uncorrected p-value = 0.01, lowest FDR corrected p-value = 0.06). Allowing for a timelag, i.e. shifting one of the microstate sequence time series relative to the other in a dyad of up to 1 second in both directions, only led to a negligible increase in the ratio of time the two participants’ microstates were synchronous (less than 1%; Supplementary Figure S18). There was also no clear consistent time-lag where inter-brain synchrony peaked, reflected by the flat line when plotting ratio of time not in the same state and the time-lag (Supplementary Figure S19).

Taken together, inter-brain synchrony, defined as cooccurrence (either simultaneous or with a constant lag) of similar microstates in both individuals, was not greater than chance level, and also not modulated by the different interaction conditions.

### 3.4 Two-brain microstate dynamics were modulated by the asymmetric mirror-game conditions

Due to the chance-level occurrence of similar global neuronal activity in the two participants in a dyad (either time-locked or time-shifted), we investigated a less constrained definition of inter-brain synchrony, by relaxing the constraint of being in the exact same microstate. This methodology is what we refer to as two-brain microstates, which tries to capture potentially different interpersonal neuronal activity states co-occurring over time in dyads.

Eight two-brain alpha microstates were determined when the microstate analysis was extended to be fitted on simultaneously recorded EEG from the 21 pairs (Figure 5A). Each two-brain microstate consists of a particular topography for one participant and a corresponding topography for the partner in the dyad at a specific time point. To distinguish them from single-brain microstates, we added a “d” (dual) prefix to their letter and a number to indicate the participant, e.g. dB1 refers to participant 1 dual-microstate B. The participant number is not so important for symmetric trials, but in the asymmetric trials we fixed the observer (in the *observer-actor* condition) or follower (in the *follower-leader* condition) to always be treated as participant 1 (top row in Figure 5A). Consequently the actor or leader were always treated as participant 2 (bottom row in Figure 5A). This was done to ensure that the effect of the asymmetric trials was not cancelled out by averaging across the asymmetry.

**Fig. 5.**
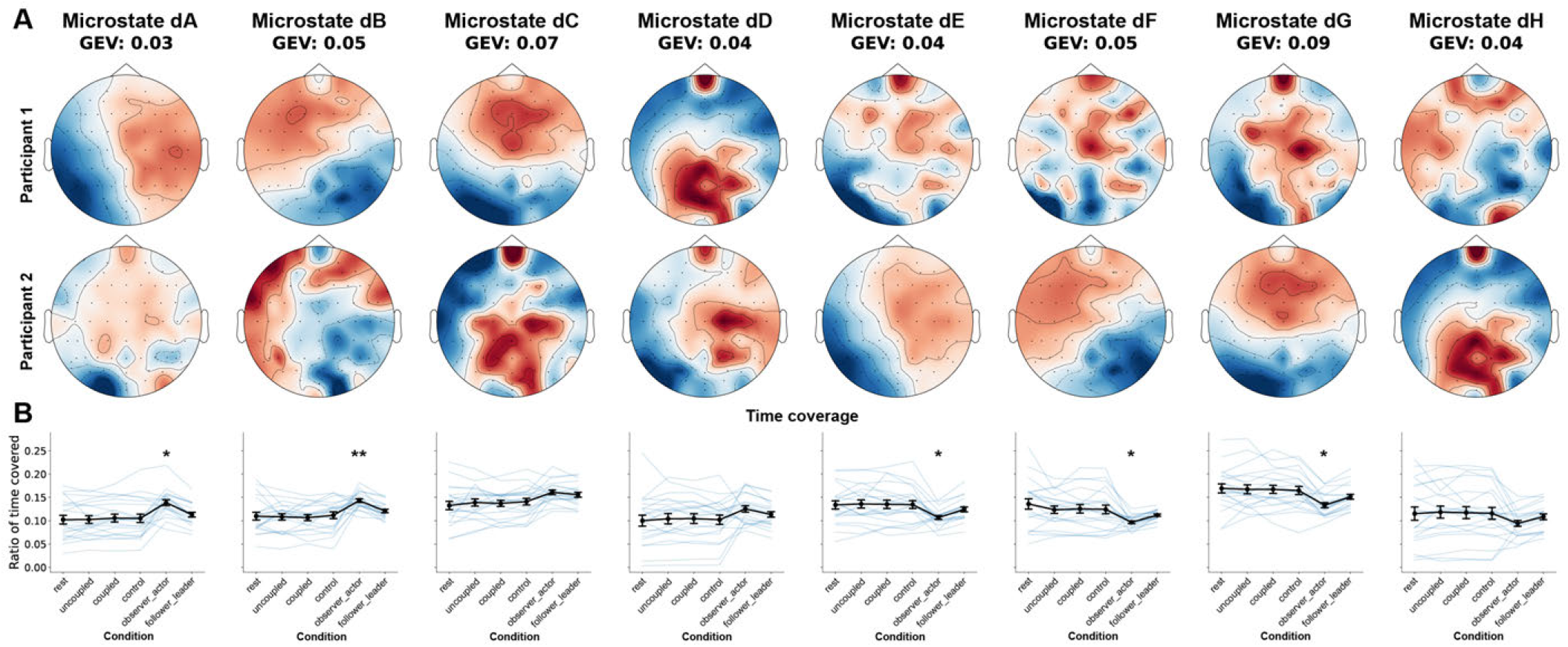
Two-brain EEG microstate analysis. A) Microstate analysis was performed on the simultaneously recorded EEG from 21 pairs collected during the mirror-game paradigm, which yielded eight two-brain alpha microstates, explaining around 41% of the variance. B) The ratio of time covered by each two-brain microstate was different between the asymmetrical and symmetrical conditions, with only the observer-actor condition differing after multiple-comparison correction. GEV: global explained variance.

Remarkably, the two-brain alpha microstates were also very similar to the conventionally found single-brain restingstate EEG microstates, with dA1, dB1, dC1 and dD1 resembling our single-brain microstates A, B, C and E and dE2, dF2, dG2, dH2 and dC2 also resembling A, B, C, E and E, respectively. The rest of the topographies were more variable and not so clearly demarcated (Supplementary Figure S20A). Further investigation of the two-brain microstate topographies revealed that the absolute values for dA2, dB2, dC2 and dD2 were very close to 0, and similarly for dE1, dF1, dG1 and dH1 (Supplementary Figure S21A), indicating that these topographic maps reflect the arbitrary average neuronal activity. In other words, if a given time point is labelled as dA, dB, dC or dD, participant 1 has a global brain activity pattern corresponding to one of the four conventionally found resting-state microstates, while participant 2 has a non-specific brain activity pattern. The opposite is true for a time point labelled as dE, dF, dG and dH, with participant 2 in one of the conventionally found resting-state microstates, while participant 1 has a non-specific activity pattern.

Interestingly, the two-brain alpha microstate dynamics showed a clear difference in the asymmetric conditions compared to the symmetric conditions (Figure 5B). When compared to the principal symmetric condition, the *coupled* interaction, we observed that dA (p-value = 0.045) and dB (p-value = 0.008) were significantly higher and dE (p-value = 0.045), dF (p-value = 0.045) and dG (p-value = 0.040) were significantly lower for the *observer-actor* condition using two-sided permutation tests with FDR multiple-comparison correction. The same direction of changes (albeit not significant after multiple-comparison correction) were also observed for the *follower-leader* condition with a higher mean in dB (uncorrected p-value = 0.06, FDR corrected p-value = 0.304), dC (uncorrected p-value = 0.036, FDR corrected p-value = 0.208) and lower mean in dF (uncorrected p-value = 0.166, FDR corrected p-value = 0.554) and dG (uncorrected p-value = 0.126, FDR corrected p-value = 0.457). Significant differences in the *observer-actor* condition after multiple-comparison correction were also found for duration (Supplementary Figure S21B; lowest FDR corrected p-value = 0.008), occurrence (Supplementary Figure S21C; lowest FDR corrected p-value = 0.008), and transition probabilities (Supplementary Figure S22; lowest FDR corrected p-value = 0.013) in the alpha frequency range.

When the two-brain microstates were fitted in the beta frequency range (Supplementary Figure S23A), the determined microstates were also similar to the conventionally determined single-brain microstates, with dA1, dB1, dC1 and dD1 and dE2, dF2, dG2, dH2 resembling our single-brain microstates A, B, C and D for each participant, respectively (Supplementary Figure S20B). However, no differences in the two-brain beta microstate features were observed for the different behavioural conditions (Supplementary Figure S23B-D and S24; lowest uncorrected p-value = 0.04, lowest FDR corrected p-value = 0.97).

Similar results were observed for two-brain microstates fitted in the broadband frequency range (Supplementary Figure S25A), with topographies resembling single-brain microstate topographies for each participant respectively (Supplementary Figure S20C), but no differences associated with the different behavioural conditions (Supplementary Figure S25B-D and S26; lowest uncorrected p-value = 0.02, lowest FDR corrected p-value = 0.83).

We also estimated two features pertaining to the complexity of the microstates: entropy and long-range temporal correlations (von Wegner et al., 2023) for two-brain microstates. No differences were observed in relation to the different mirror-game conditions for entropy, in any of the three frequency ranges (Supplementary Figure S27A-C; lowest uncorrected p-value = 0.34, lowest FDR corrected p-value = 0.93). There were significantly more long-range temporal correlations in the *rest* condition compared to the principal *coupled* condition, for all three frequency ranges, while the *uncoupled* condition also had higher long-range temporal correlations in the beta and broadband frequency band and *control* and *observer-actor* condition had higher long-range temporal correlations than the *coupled* in the beta frequency band (Supplementary Figure S27D-F; all FDR corrected pvalues < 0.01).

In order to discern whether the changes in two-brain alpha microstate dynamics were specific to the interaction itself, as opposed to an effect of the asymmetric behavioural assignment, we computed the microstate features for pseudo-pairs. We observed that the pseudo-pairs also showed a significant difference in two-brain microstate dynamics for the asymmetric conditions, and there was no difference in time coverage between the real pairs and the pseudo-pairs (Supplementary Figure S28A; lowest uncorrected p-value = 0.734, lowest FDR corrected p-value = 0.994). The lack of difference was also observed in duration, occurrence and transition probabilities (lowest uncorrected p-value = 0.349, lowest FDR corrected p-value = 0.997) suggesting that the changes in two-brain alpha microstate dynamics were not specific to the interaction, but rather the asymmetric task.

There were also no differences in entropy between real and pseudo-pairs (Supplementary Figure S28B; lowest uncorrected p-value = 0.771, lowest FDR corrected p-value = 0.900) and long-range temporal correlations between real and pseudo-pairs (Supplementary Figure S28C; lowest uncorrected p-value = 0.392, lowest FDR corrected p-value = 0.796).

Overall, we observed that microstate analysis applied to dyads in the mirror-game yielded two-brain microstates corresponding to the canonical single-brain microstates in one of the participants, while the other participant was in an arbitrary non-specific neural activity state. However, while single-brain microstate dynamics were not related to the different mirror-game conditions, two-brain microstates revealed that the asymmetric conditions were associated with more asymmetric joint spatiotemporal dynamics, with the observer being in the canonical single-brain resting-state microstates to a greater degree than the actor. This modulation was specific to the asymmetric task itself, rather than the reciprocal social interaction.

### 3.5 Source localization of two-brain microstates

To determine the underlying cortical neural sources for each two-brain microstate, we applied eLORETA to the microstate topographies. Due to the high similarities between the topographies obtained from the three frequency ranges, only the two-brain alpha microstates are shown (Figure 6). We observed that the cortical neural sources corresponding to the four conventionally found resting-state microstates were similar between the two participants in a pair. Focusing on the areas with greatest activity, we observed that dA1 and dE2 showed high activity in left temporal and parietal areas and low activity in left temporal and right cingulate and parietal areas (Figure 7A and 7E). dB1 and dF2 showed high activity in right parietal, temporal and cingulate areas and low activity in right temporal and left cingulate areas (Figure 7B and 7F). dC1 and dG2 showed high activity in left cingulate and temporal areas and low activity in Left and right temporal areas (Figure 7C and 7G). dD1 and dH2 showed high activity in left and right frontal and left anterior cingulate areas and low activity in left occipital areas (Figure 7D and 7H). Notice that due to the microstate polarity invariance during the clustering, the high and low activity labels are arbitrary and interchangeable. The keypoint is that there is a difference in potential between the areas listed in high and low. Additionally, the microstates reflecting the arbitrary mean neuronal activity (dA2, dB2, dC2, dD2, dE1, dF1, dG1 and dH1) were omitted as their corresponding source activities were also around 0. For readers unfamiliar with flatmaps, Supplementary Figure S29 shows the underlying sources with the more standard view of the cortical regions on the inflated brain.

**Fig. 6.**
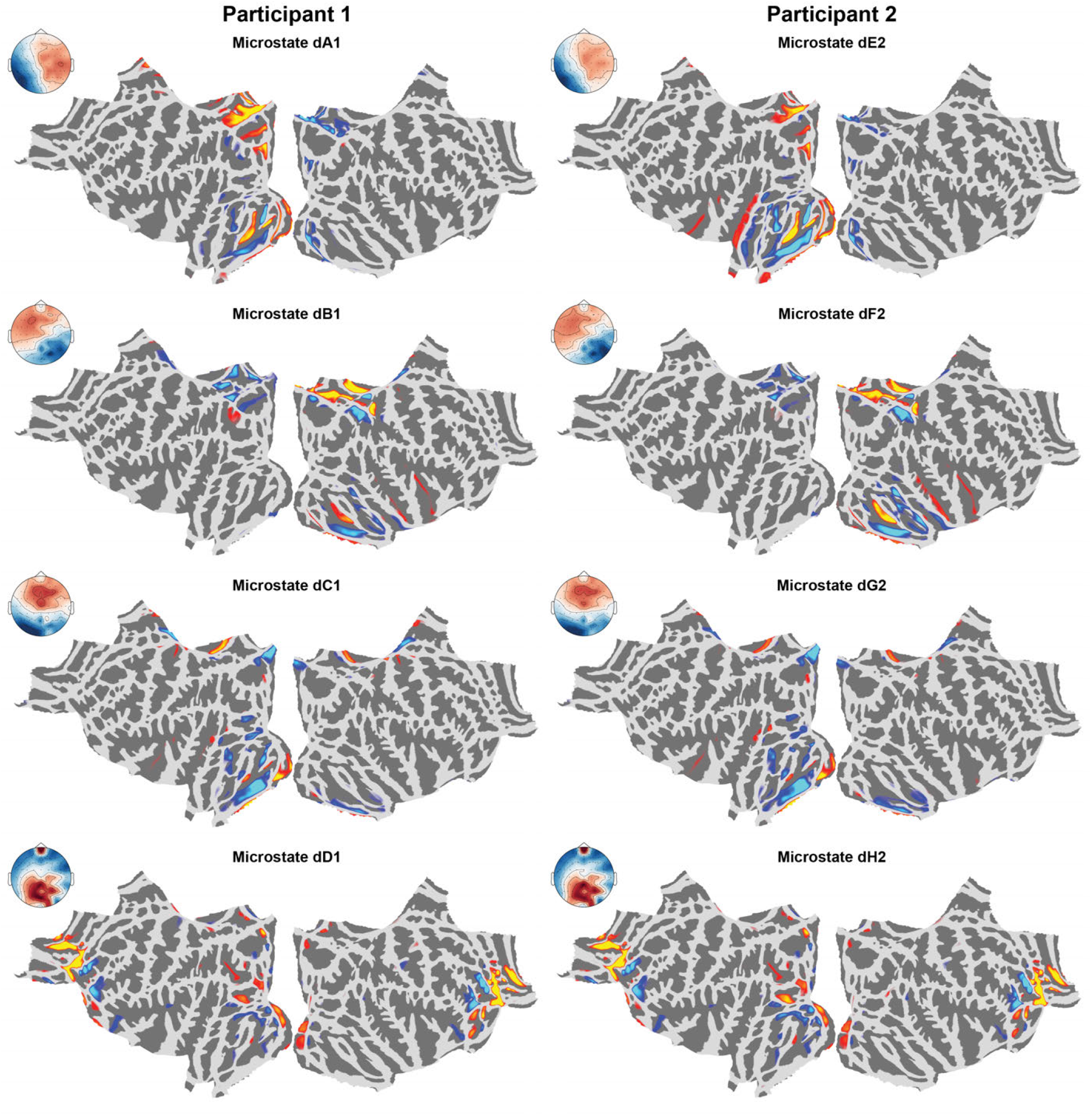
The eight two-brain alpha microstates and their corresponding eLORETA cortical activities. We only show the cortical sources for dA1, dB1, dC1, dD1, dE2, dF2, dG2 and dH2, as the other two-brain microstates reflected the mean arbitrary activity. See Supplementary Figure S29 for the same data plotted with an inflated brain view.

**Fig. 7.**
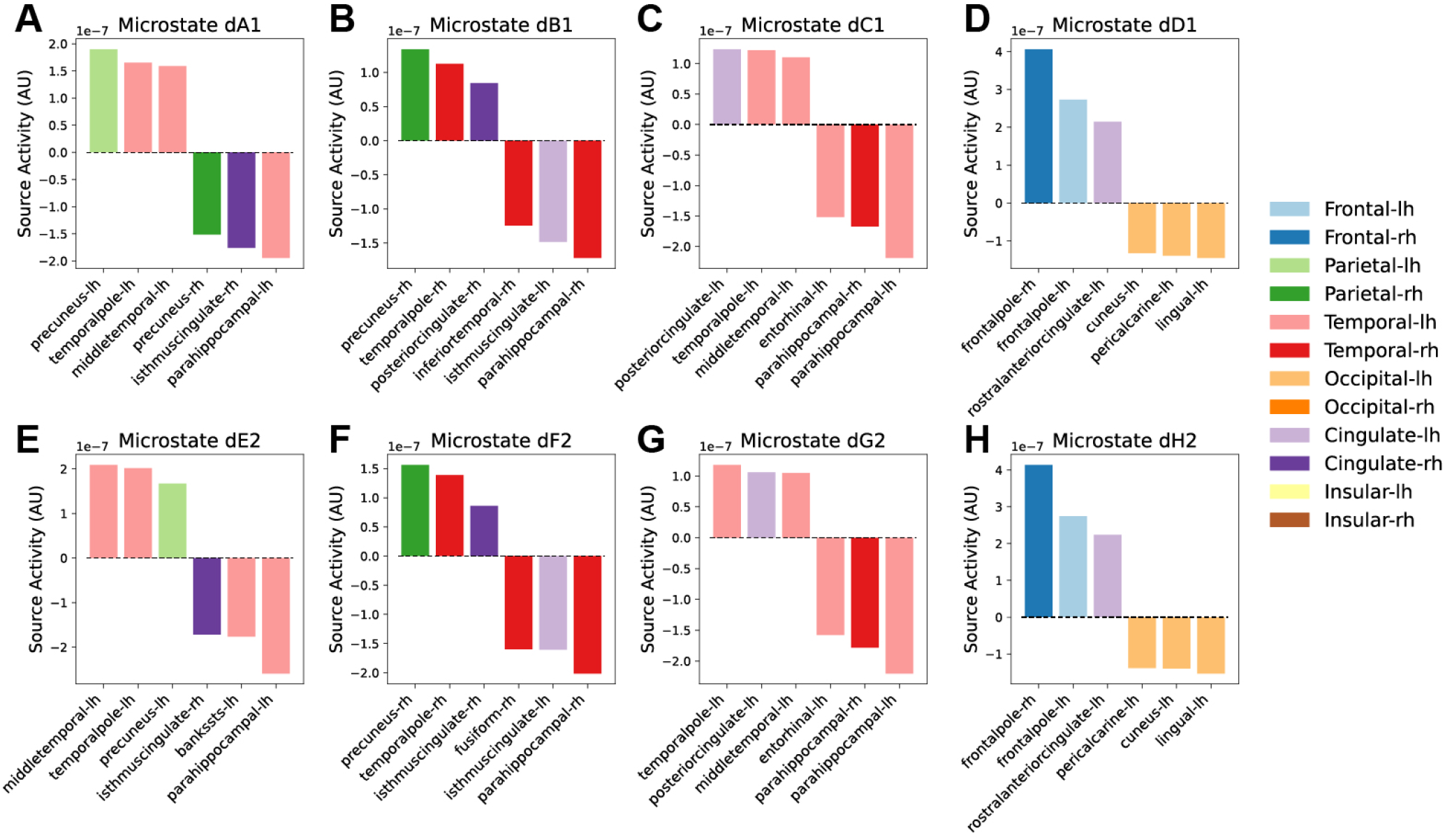
Top three positive and negative areas in the eight source localized two-brain alpha microstates. The colors indicate which brain region the corresponding area belong to, while the shade indicate whether it is the left or high hemisphere, with darker colors belonging to the right hemisphere. The dashed black line indicates zero activity. Notice that due to the microstate polarity invariance during clustering, the positive or negative activity signs are interchangeable.

In summary, the two-brain microstates were primarily generated by neural activity in the following cortical brain areas: dA1/dE2 from left temporal/parietal and posterior cingulate areas; dB1/dF2 from right temporal/parietal and posterior cingulate areas; dC1/dG2 from left temporal and posterior cingulate areas; and dD1/dH2 from frontal, anterior cingulate and left occipital areas.

## 4 Discussion

In this study, we developed a novel method for investigating inter-brain synchrony and inter-brain asymmetry in a hyperscanning EEG experiment. Specifically, we expanded the microstate analysis framework to dyads of interacting participants, enabling us to investigate quasi-stable moments of synchronous, as well as asymmetric simultaneously occurring spatiotemporal activity. We found that conventional microstates fitted to individuals (single-brain microstates) were not related to the different task conditions; however, the dynamics of the two-brain alpha microstates were changed for the *observer-actor* condition, compared to all other conditions where participants had more symmetric task demands (rest, individual, joint). Interestingly, the topographies of the two-brain microstates were related to the conventionally found resting-state microstates determined from single individuals (Tarailis et al., 2023), and our source localized two-brain microstates (Figure 6) also had cortical activities similar to previous findings relating the microstates to the Default Mode Network (DMN) (Pascual-Marqui et al., 2014).

One often discussed topic regarding microstate analysis is the frequency range of the data the clustering should be fitted to. In the early microstate papers, the alpha frequency range was employed (Lehmann et al., 1987), but later papers found that the resting-state microstates could also be determined using broadband, e.g. 2-20 Hz or 1-40 Hz, which has subsequently become the more standard approach (Tarailis et al., 2023). However, alpha oscillations are regarded as the main driving component for broadband microstates (Milz et al., 2017; von Wegner et al., 2021). As our EEG data were not only from a resting-state condition, but recorded during an interactive mirror-game paradigm where the participants were producing motor movements, we were also interested in the beta frequency range (13-30Hz), which is known to be important for motor movements (Conway et al., 1995; Perez et al., 2006; Baker, 2007). A recent paper (Férat et al., 2022) also demonstrated that despite the near-perfect spatial correlations between microstate topographies across frequencies for both eyes open and eyes closed at rest, spectrally specific microstate analysis yielded independent information about spatial-temporal dynamics, with alpha-band microstates classifying eyes open from eyes closed EEG better than broadband microstate features (Férat et al., 2022). Therefore, we performed the microstate analysis for all three frequency ranges of interest in parallel. Consistent with previous results (Férat et al., 2022) the topographies of our determined singlebrain microstates were very similar across frequency ranges, and we also showed that two-brain microstates were similar across frequencies. However, we only found significant differences associated with the asymmetric *observer-actor* condition in the alpha frequency range for the two-brain microstates. Previous studies have also found associations between alpha/mu oscillations and motor control, joint attention, and coordination (Klimesch et al., 2007; Tognoli et al., 2007; Lachat et al., 2012; Dumas et al., 2012), e.g., showing role-specific modulation of alpha-band activity when participants lead or follow (Konvalinka et al., 2014; Flösch et al., 2023) or when different interpersonal strategies emerge (Heggli et al., 2021). Our findings showing that two-brain alpha microstates were specifically associated with differences during the asymmetric tasks is also consistent with the recent hyperscanning dual EEG findings by Flösch et al. (2023), who found differences in alpha and low beta oscillations over large areas (frontal, central and parietal areas) in dyads performing a cooperative task with asymmetric roles (sender and receiver).

The functional role of our findings is still unclear and further investigation is needed to better understand the neural mechanisms. However, the cortical areas associated with our determined microstates have been postulated to be a fragmented high time resolution version of the slow metabolically (PET/fMRI) determined Default Mode Network (DMN) (Pascual-Marqui et al., 2014). Given the similar trends with higher presence of dA, dB, dC and dD in the observer and lower presence of dE, dF, dG and dH in the actor, we hypothesize that the effect we observe is due to the observer being in a more resting-state DMN-like neuronal brain state compared to the actor. Importantly, this effect is only observed for the two-brain microstates and not found for single-brain microstates, indicating that both participants in the dyads have DMN activity; however, the relative degree of activity might not be equal, and this difference can only be determined when analysing both participants simultaneously. An important point about two-brain microstates is that one time point is always associated with only one label, hence if the label is dA, then it means participant 1 is more in a restingstate microstate A than participant 2. They are competing for label assignment, whereas in single-brain microstates they both get assigned a microstate for each time step, thus there is no competition or comparison between the individuals. The difference we observed between the symmetric interactive *coupled* condition and *observer-actor* condition could thus correspond to the DMN being more active in the observer relative to the actor, compared to the symmetrical trials where the DMNs are equally active.

Another discussed topic regarding microstate analysis and unsupervised clustering is the number of clusters that should be extracted. Too few might not capture enough information about the data; however, too many clusters could lead to overfitting, which result in the clusters not being able to generalize. There are various methods to determine an optimal number of microstates and it is also possible to use several methods in combination, e.g. the meta-criterion (Custo et al., 2017). We used the cross-validation criterion (PascualMarqui et al., 1995) and found five and eight microstates to be optimal for the single-brain and two-brain analysis, respectively. One might argue that one reason for not finding any task-related differences in single-brain microstates could be due to the lower number of extracted microstates, and subsequently lesser information captured. However, the fact that the topographies of the two-brain microstates show high similarity to four of the single-brain microstates, provides some evidence that they are not capturing less information. Additionally, the global explained variance was higher for the single-brain microstates compared to two-brain microstates, most likely due to the concatenated two-brain EEG time-series being more variable and thus harder to reduce into specific quasi-stable states. To fully resolve this issue, we also repeated the single-brain microstate analysis with eight clusters and we still did not observe any task-related differences in microstate dynamics (Supplementary Figure S30). Noticeably, the global explained variance only increased with 4% by adding three more microstates, emphasizing the point that once too many clusters are added, the clusters might begin to overfit and only fit to smaller (potentially noisy) portions of the data.

The oscillatory nature of neural activity of EEG is also one of the reasons why the *K*-means algorithm had to be modified to be polarity invariant for the determination of microstates (Pascual-Marqui et al., 1995). One of the challenges of expanding the microstate analysis from single individuals to dyads was to retain the polarity invariance for both individuals in a pair. The standard implementation to ignore polarity in microstates is to square the spatial correlation coefficient (von Wegner et al., 2021). But this solution does not ensure that the individual topographies from each participant in a two-brain microstate were polarity invariant. For example in the case of dA1 and dA2: if we define them as having a positive polarity, then the topographies can be written as {+, +}. If the microstates are polarity invariant, the following polarities should be equivalent: {+, +}, {+, -}, {-, +} and {-, -}. However, using the squared spatial correlation coefficient only ensures {+, +} will be treated the same as {-, -}, but a neural activity pattern similar to {+, -} or {-, +} would not be equivalent. Another method to make microstates polarity invariant is to use the absolute transformation (Tait and Zhang, 2022). Initially, we also tried the absolute transformation, and while the microstates were polarity invariant, there is a large caveat with this method, namely that negative and positive values will be treated equally, and this fundamentally changes the data, e.g. if Fp1 is 20µV, Fp2 is 16µV, O1 is -19µV and O2 is -21µV, then after the absolute transformation Fp1 would be seen as more similar to O1 and O2 as opposed to Fp2. Such a dramatic change of the data would also change what kind of microstates will be determined. Ultimately, we decided to compute the spatial correlation for each time point with all four combinations of possible polarity configurations, and chose to keep the highest correlation, to preserve the polarity invariance for the microstate determination, without compromising the input data.

Remarkably, both the single-brain and two-brain microstate topographies were similar to the commonly found resting-state EEG microstates in previous literature (Michel and Koenig, 2018; Tarailis et al., 2023), despite being extracted from the motor coordination tasks during the mirror-game. One reason for this similarity could have been due to the presence of the 2min *rest* condition at the start and end of the experiment. To address this, we also tried fitting the two-brain microstates without the *rest* conditions and observed similar two-brain microstates (Supplementary Figure S31), indicating that the reason for the similarity to the commonly found resting-state single-brain EEG microstates was not due to being fitted on *rest* conditions.

It has not escaped our notice that we were not able to find inter-brain synchrony, either as simultaneous occurrence of inter-brain microstates, or as quasi-stable co-occurring brainstates in the dyads. In fact, the amount of time the two participants were in the same microstate was not different from chance-level (Supplementary Figure S17). However, our two-brain microstates method does not capture functional connections between brains, hence it is not directly comparable to more conventional IBS analyses employing phase or amplitude coupling between brains. It is thus free from any assumptions regarding the existence of coupling between two persons’ brain signals. Instead, it captures whether people are likely to simultaneously be in the same (symmetric) or asymmetric spatiotemporal states at the same time. In addition, the determined two-brain microstates, which are optimized to explain as much of the variance in the data as possible, all corresponded to a stable canonical microstate in one participant, while the other participant had activity close to 0 and hence corresponded to arbitrary undetermined activity. This does not mean that there was not any inter-brain synchrony during behavioural synchronization or interaction, as the microstates only focus on global neuronal brain states, hence smaller spatially focal synchronous areas would be overlooked by this methodology. It would be interesting for future work to investigate comparisons between focal interbrain synchronization (e.g., utilizing phase locking value or circular correlations (Xu et al., 2024; Zamm et al., 2024)) and global brain state synchrony using microstates.

Shared neuronal spatiotemporal responses across brains have also previously been investigated using inter-subject correlations (ISC) (Nastase et al., 2019)). ISC based methods investigate the shared information across subjects exposed to the same time-locked stimuli, e.g., watching the same video clips, and can be applied without hyperscanning (Dmochowski et al., 2012; Nastase et al., 2019). Conceptually, some parallels can be drawn between correlated component analysis (CorrCA) (Dmochowski et al., 2012; Parra et al., 2019)), an ISC based EEG method, and our two-brain microstates. Both of these methods extract shared decompositions, e.g. in the form of common topographical maps, across subjects; however, CorrCA does this by maximizing ISC over time and across subjects to obtain shared neuronal spatiotemporal responses (Parra et al., 2019), whereas our two-brain microstates were determined based on the amount of global explained variance over time across dyads. This difference means our method is not constrained to only investigate shared symmetric spatiotemporal dynamics, but also potentially co-occurring asymmetric spatiotemporal dynamics. The potential to investigate simultaneously occurring asymmetric whole-brain neural dynamics, on a fine-grained temporal scale, has a broad range of applications for investigating social interactive and joint action paradigms, where actors might have different complementary roles (Sebanz et al., 2006; Masumoto and Inui, 2013; Skewes et al., 2015; Sartori and Betti, 2015; Sebanz and Knoblich, 2021; Konvalinka et al., 2023).

## 5 Conclusion

Taken together, two-brain microstates might serve as a novel method for identifying both synchronous and asymmetric spatiotemporal inter-brain states during real-time social interaction. In this study, we show that two-brain microstates are modulated by asymmetric interactions, which are not driven by the interaction itself but by the asymmetry in the tasks. To further validate the method, future work should apply the methodology to other behavioural tasks where asymmetric behaviour is both intentional (instructed) as well as spontaneous. In addition, the methodology has an added benefit of being expandable, thus multi-brain microstates could be utilized to investigate neural mechanisms underlying social interaction in groups of interacting individuals.

## Data and code availability

The raw EEG data that support the findings of this study are openly available (Zimmermann et al., 2022) from Figshare at https://figshare.com/s/5e3dcb56ed515c2cb786. The code used for this study is available from https://lab.compute.dtu.dk/ glia/two-brain-eeg-analysis.

## CRediT authorship contribution statement

Q.L.: conceptualization, data curation, formal analysis, methodology, project administration, software, visualization, writing-original draft and writing-review and editing; M.Z.: data curation, investigation, software, writing-review and editing; I.K.: conceptualization, funding acquisition, investigation, resources, supervision, writing—original draft and writing—review and editing. All authors approved the final version of the manuscript.

## Acknowledgements

This work is supported by the Villum Young Investigator grant (project no. 37525).

## Supplementary Materials

### Supplementary Methods

#### Mirror-game instructions and details

Participants were instructed to produce free movements either alone (*uncoupled*) or synchronized without a designated leader (*coupled*). In the *follower-leader* condition, the leader was instructed to produce free movements, while the follower was instructed to imitate (mirror) these movements. For the *observer-actor* condition, the actor was instructed to produce free movements, while the observer was instructed to observe. Finally, in the *control* condition both participants were asked to imitate (mirror) the dot movements on the screen, which was the same (but mirrored) for both participants hence a control for the similarity in movements produced in the coupled condition, without the interpersonal interaction. Each trial was preceded by a short label informing about the upcoming condition, and, where applicable, the role of each participant. Trial order was pseudo-randomized, such that each block of five trials contained all experimental conditions, and conditions did not repeat in consecutive trials. For the *observer-actor* and *follower-leader* conditions, the assigned roles were alternated between participants. See Zimmermann et al. (2022) for more details.

#### Detrended fluctuation analysis in two-brain microstates

To perform the DFA analysis, the microstate sequence time series had to be partitioned into two classes (Van De Ville et al., 2010), and given the asymmetric topographies with dA, dB, dC and dD reflecting participant 1 being in a conventional resting-state microstate, and dE, dF, dG and dH reflecting participant 2, respectively, we partitioned the two classes as *C*_1_ = {*dA, dB, dC, dD*}, *C*_2_ = {*dE, dF, dG, dH*}.

## Supplementary Figures

**Fig. S1.**
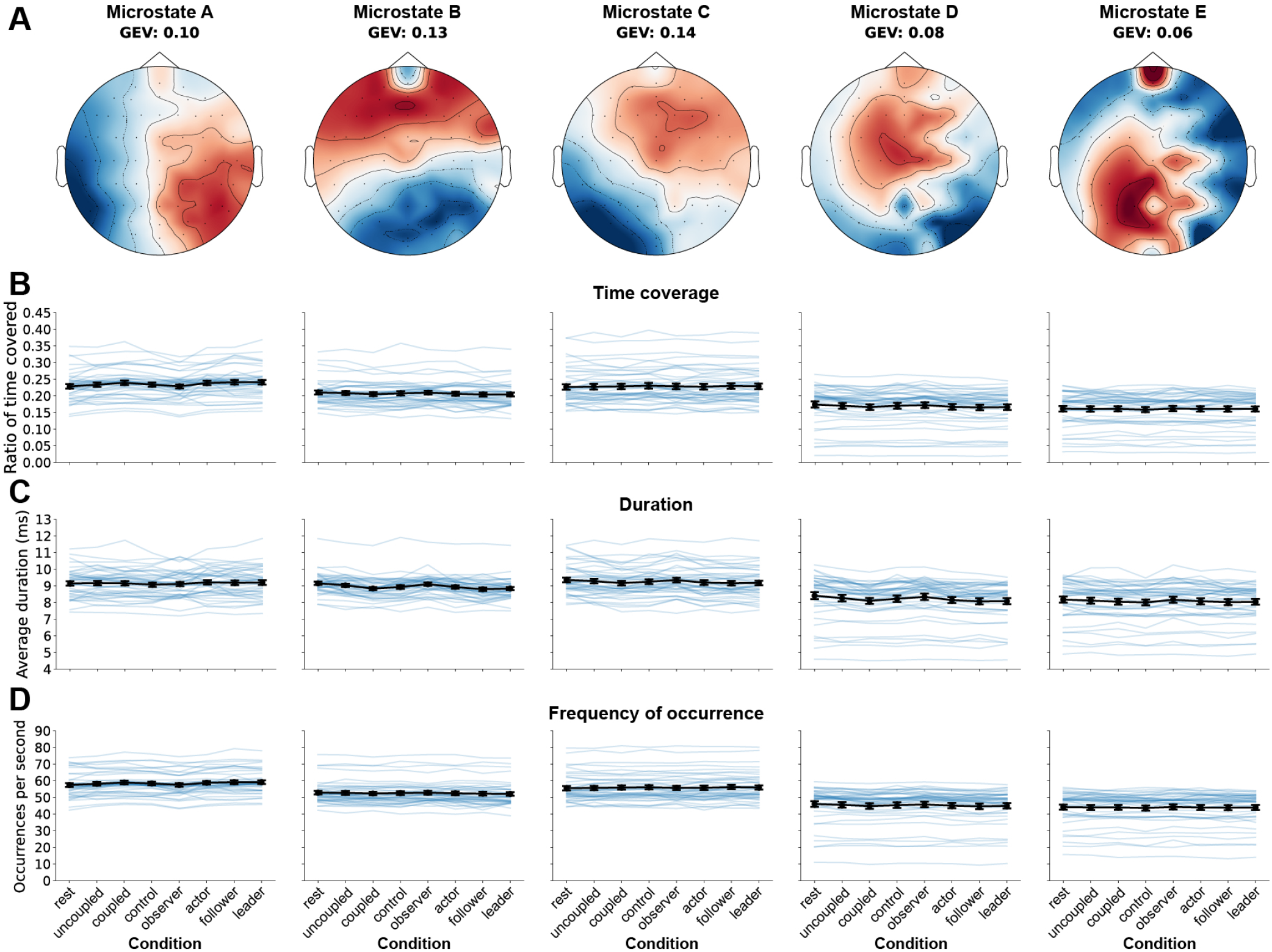
Conventional single-brain EEG beta microstate analysis. A) Microstate analysis was performed on the 42 individual EEG timeseries filtered in the beta frequency range and five microstates were determined, which explained around 50% of the variance. B) The ratio of time covered of each microstate. C) The average duration of each microstate. D) The frequency of occurrence for each microstate. None of the features were related to the different behavioural conditions. GEV: global explained variance.

**Fig. S2.**
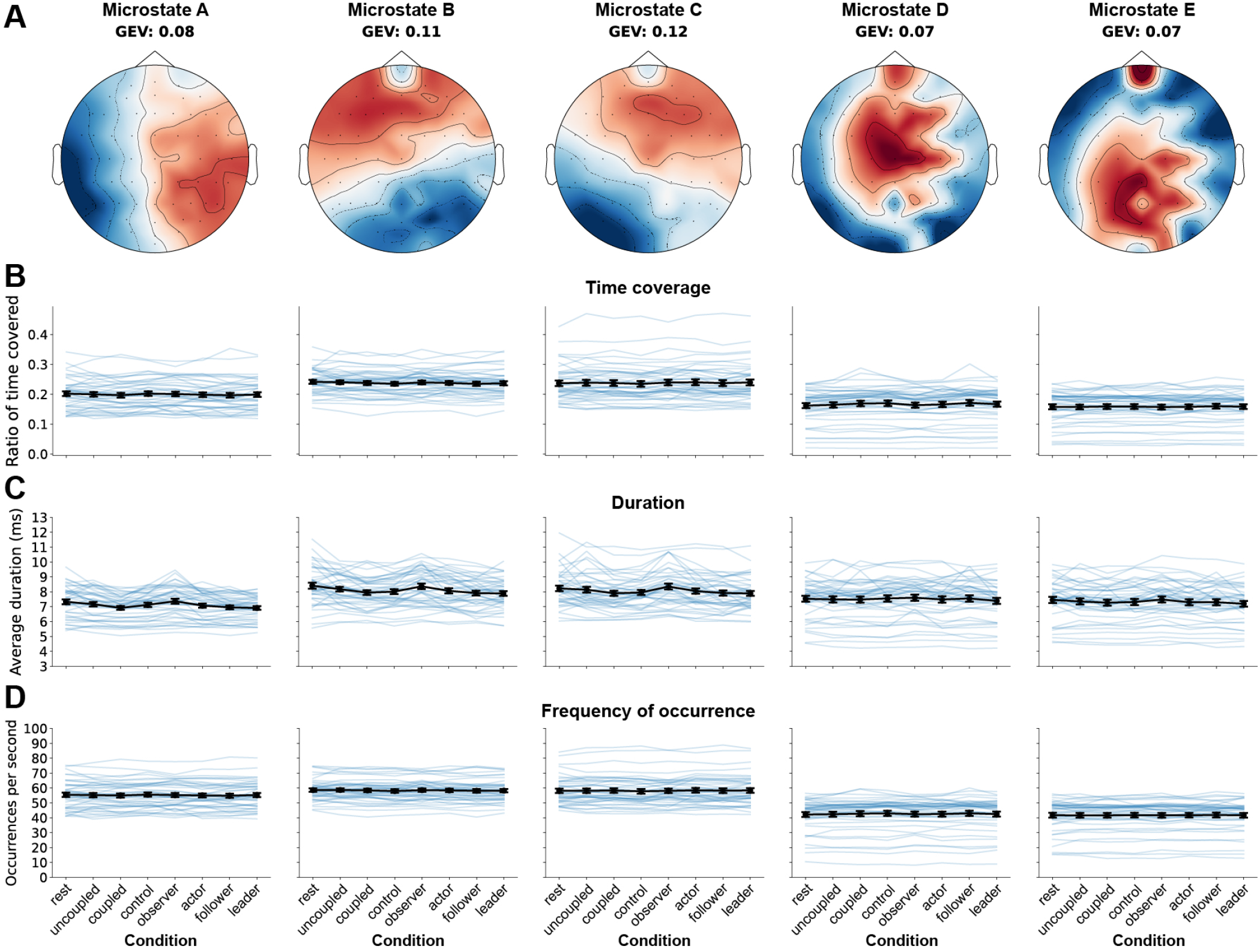
Conventional single-brain EEG broadband microstate analysis. A) Microstate analysis was performed on the 42 individual EEG timeseries filtered in the broadband frequency range and five microstates were determined, which explained around 44% of the variance. B) The ratio of time covered of each microstate. C) The average duration of each microstate. D) The frequency of occurrence for each microstate. None of the features were related to the different behavioural conditions. GEV: global explained variance.

**Fig. S3.**
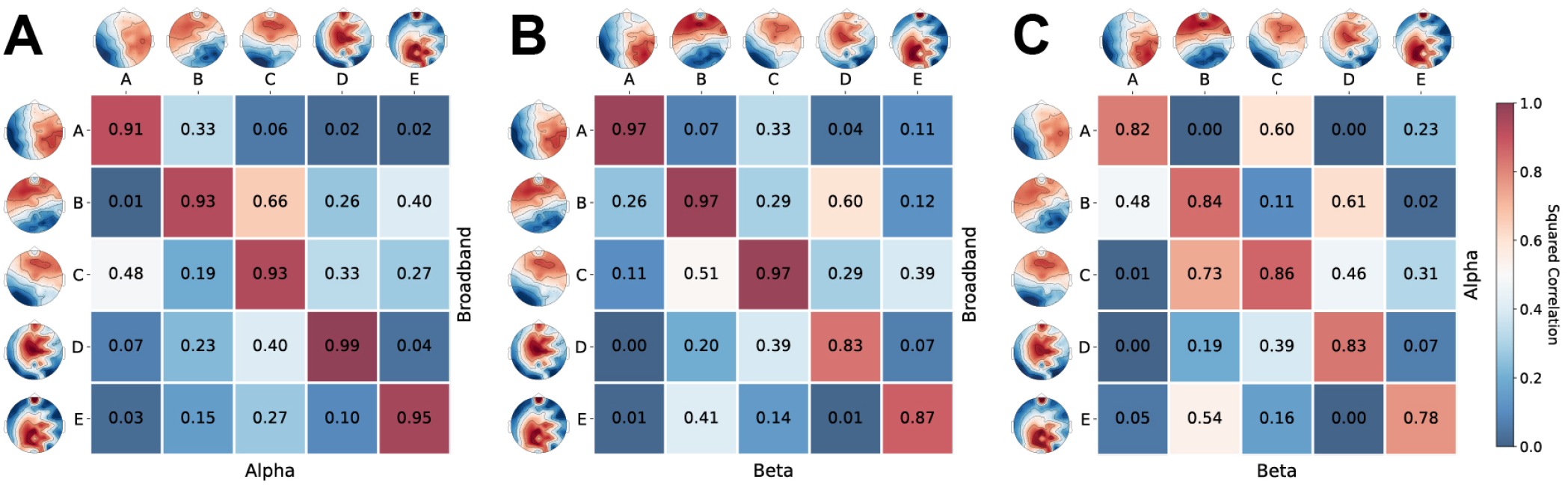
Spatial correlation between single-brain microstate topographies across frequencies. The squared correlations between A) broadband and alpha, B) broadband and beta, and C) alpha and beta frequency ranges.

**Fig. S4.**
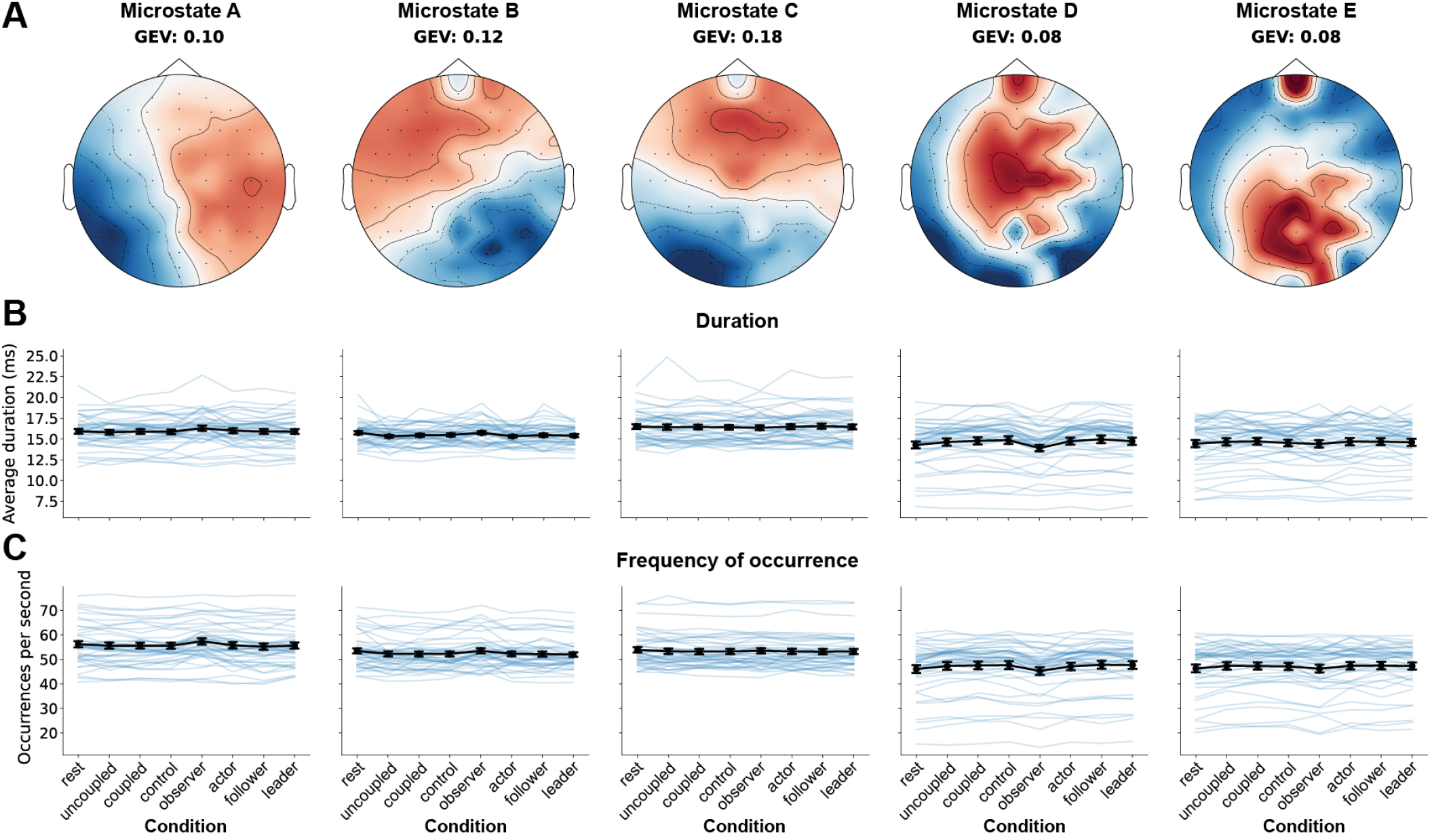
Conventional single-brain EEG alpha microstate analysis. A) Microstate analysis was performed on the 42 individual EEG timeseries filtered in the alpha frequency range and five microstates were determined, which explained around 56% of the variance. B) The average duration of each microstate. C) The frequency of occurrence for each microstate. None of the features were related to the different behavioural conditions. GEV: global explained variance.

**Fig. S5.**
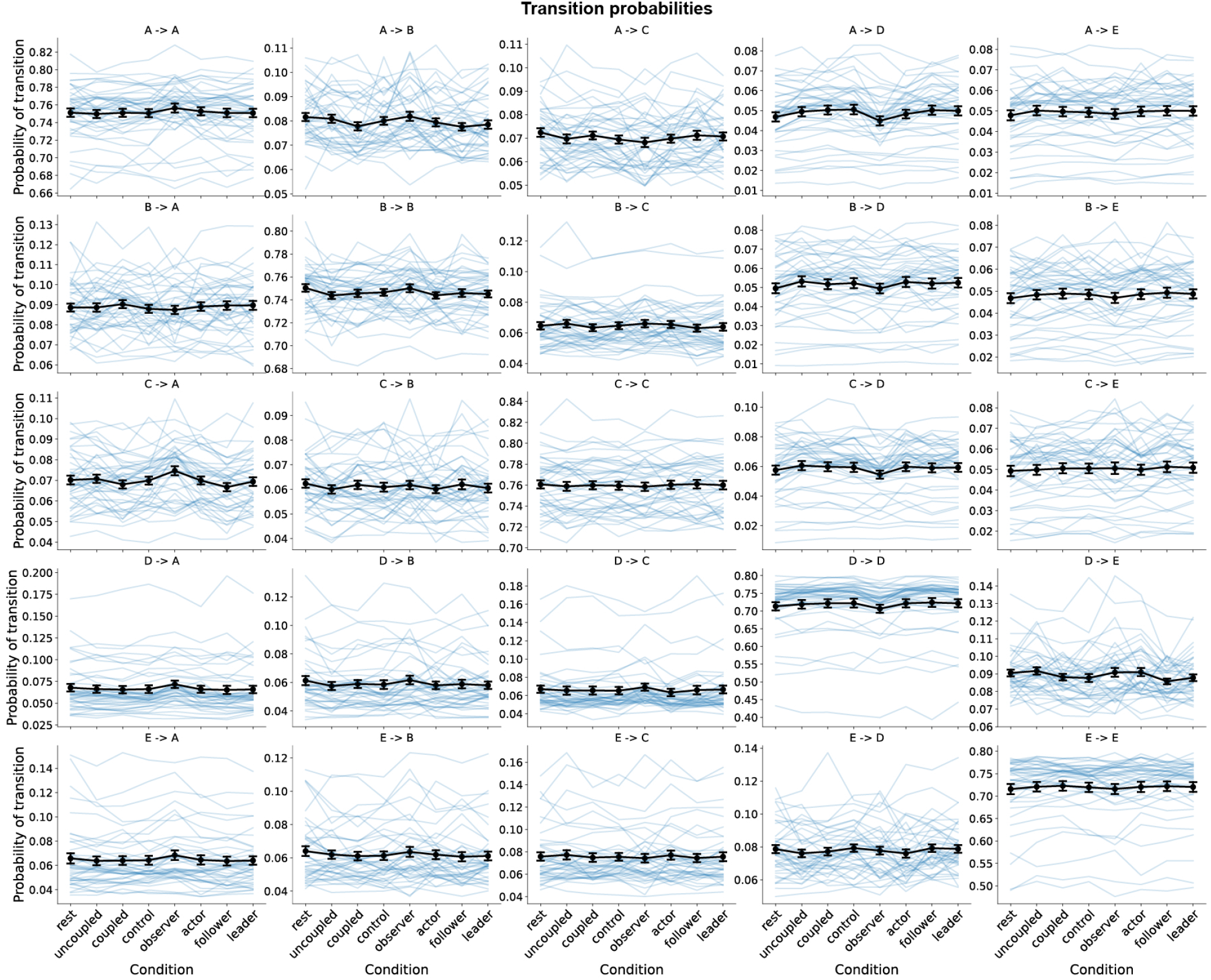
Single-brain alpha microstate transition probabilities. The transition probabilities for single-brain alpha microstates. None of the features were related to the different behavioural conditions.

**Fig. S6.**
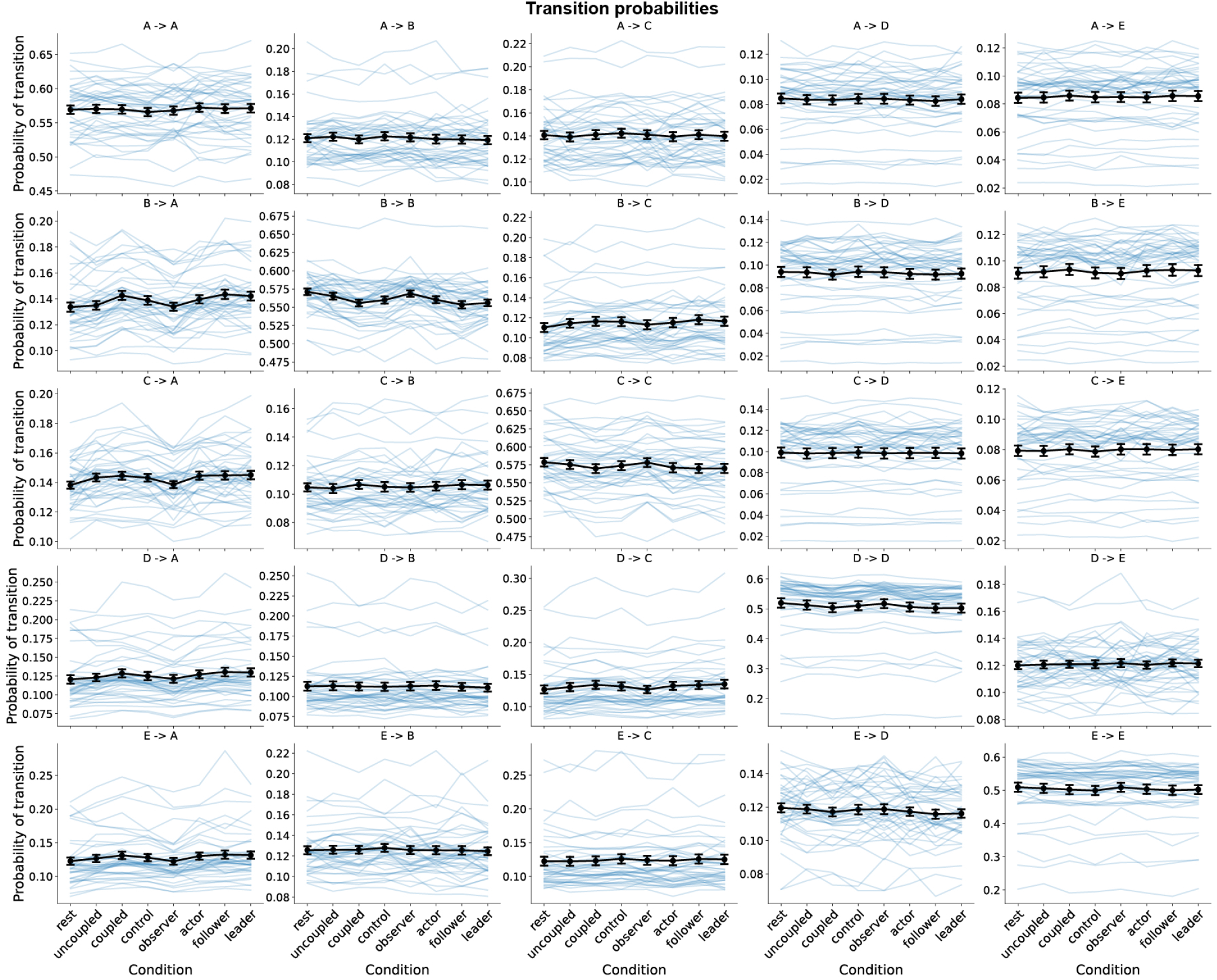
Single-brain beta microstate transition probabilities. The transition probabilities for single-brain beta microstates. None of the features were related to the different behavioural conditions.

**Fig. S7.**
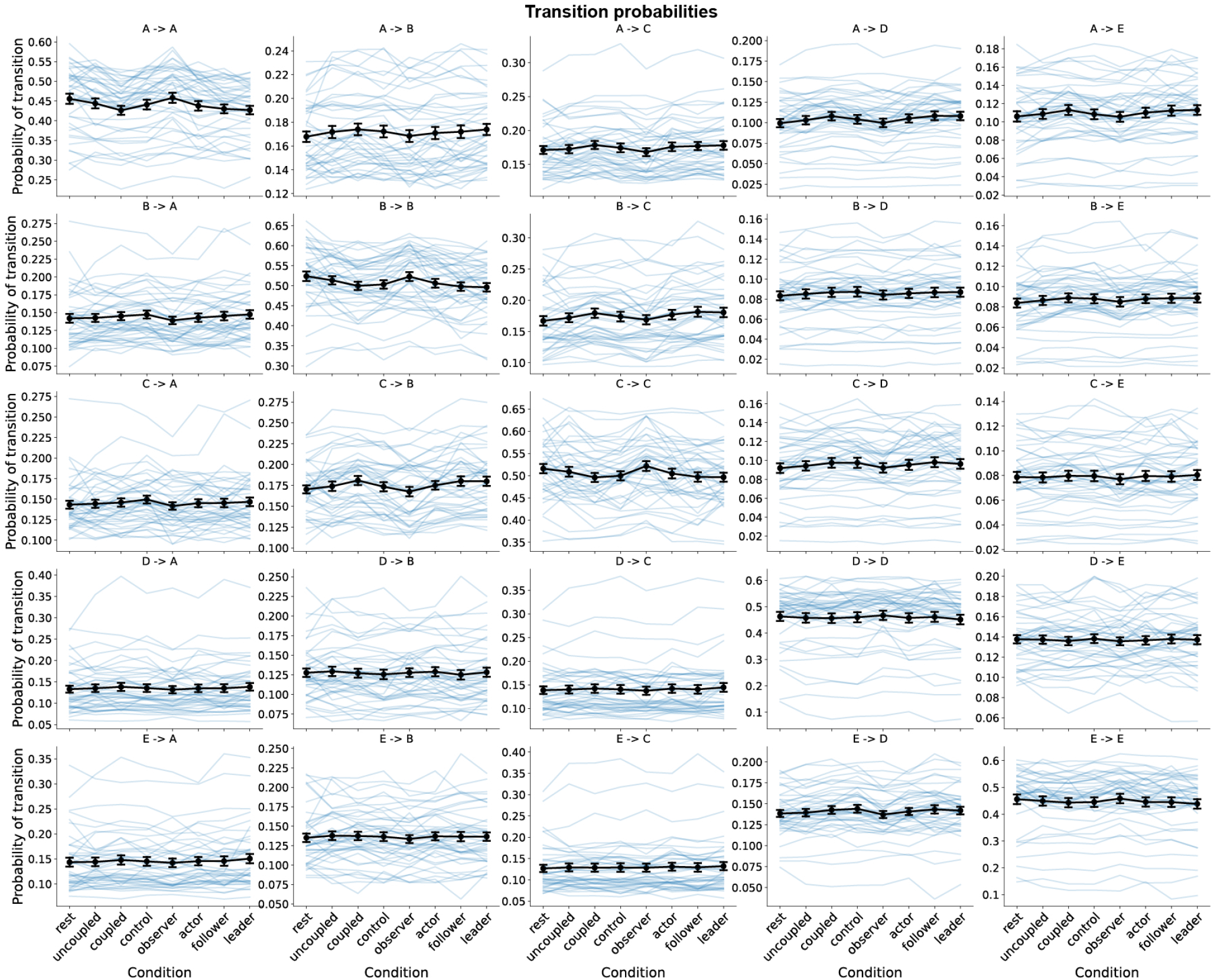
Single-brain broadband microstate transition probabilities. The transition probabilities for single-brain broadband microstates. None of the features were related to the different behavioural conditions.

**Fig. S8.**
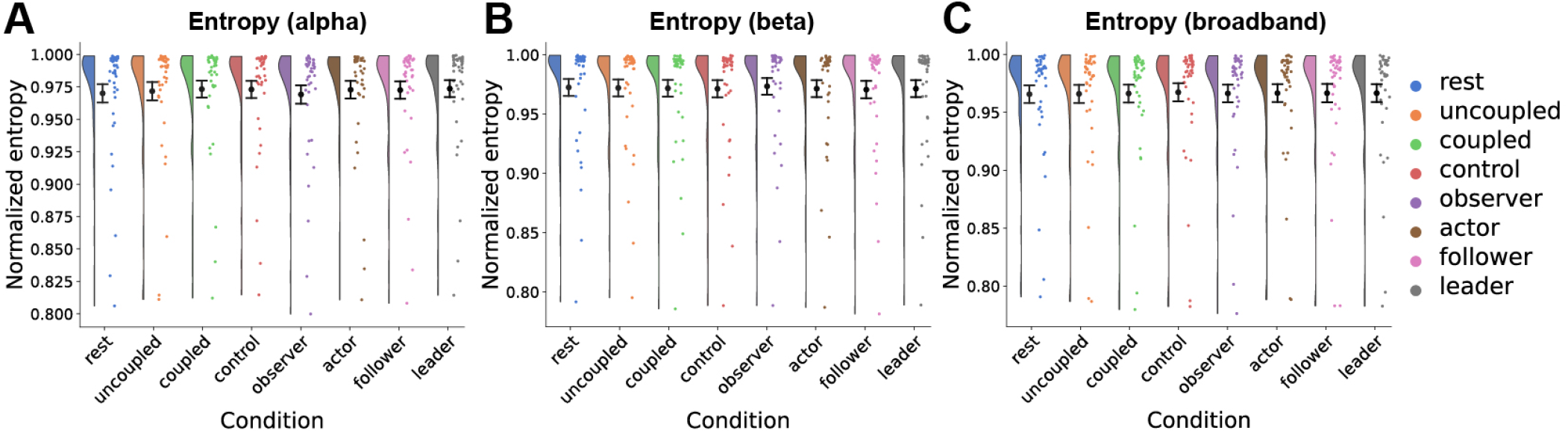
Single-brain EEG microstate entropy. Shannon entropy was computed for the microstate sequence time series determined in the A) alpha, B) beta and C) broadband frequency range, normalized to the theoretical maximum (uniform label distribution). Entropy were not related to the different behavioural conditions.

**Fig. S9.**
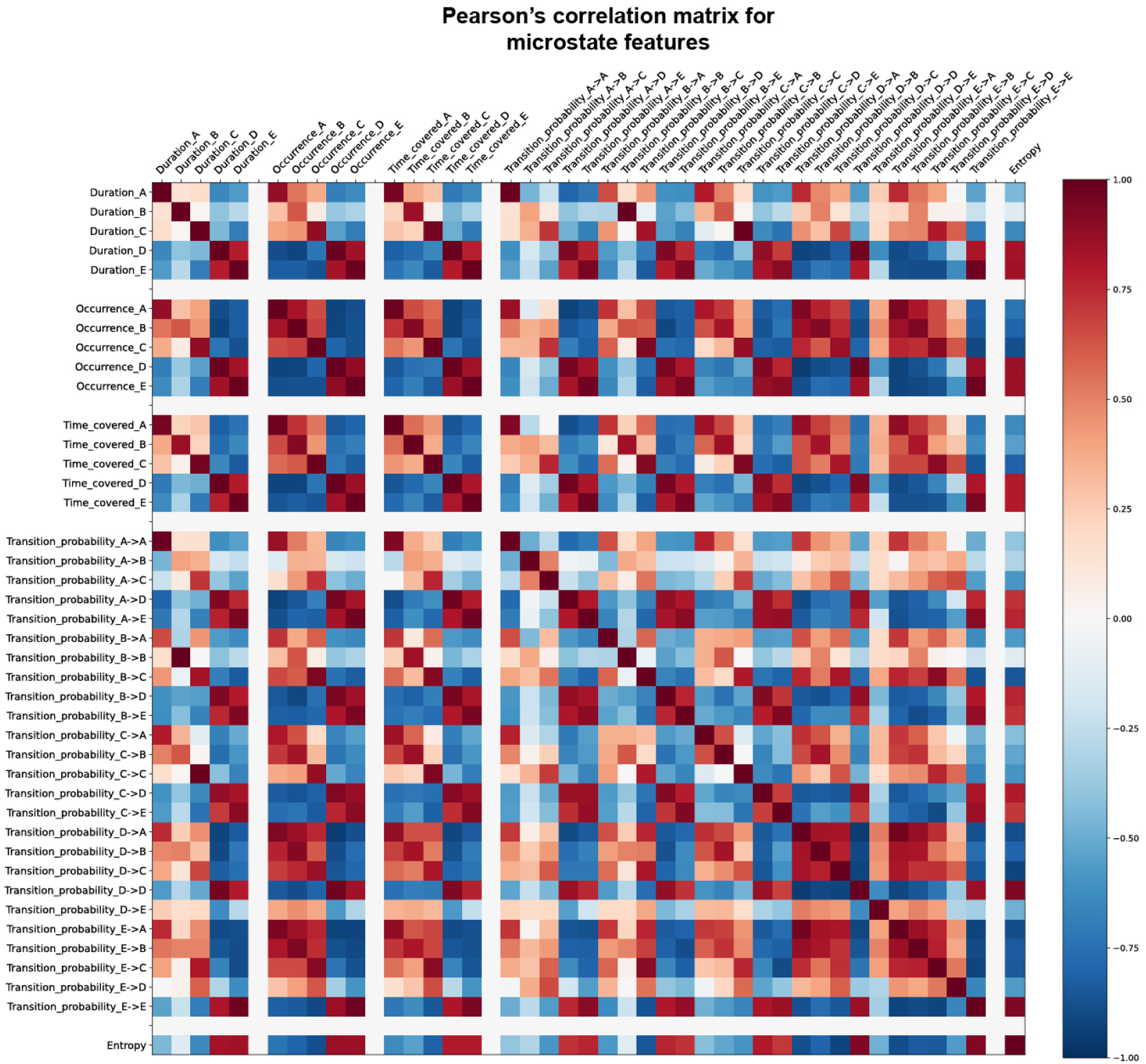
Pearson’s correlation for single-brain EEG alpha microstate features. Strong positive correlations are seen between microstate features corresponding to the same microstate. Microstate A, B and C are also often positively correlated, and the same is true for D, E, due to the high similarity in their corresponding topographic maps.

**Fig. S10.**
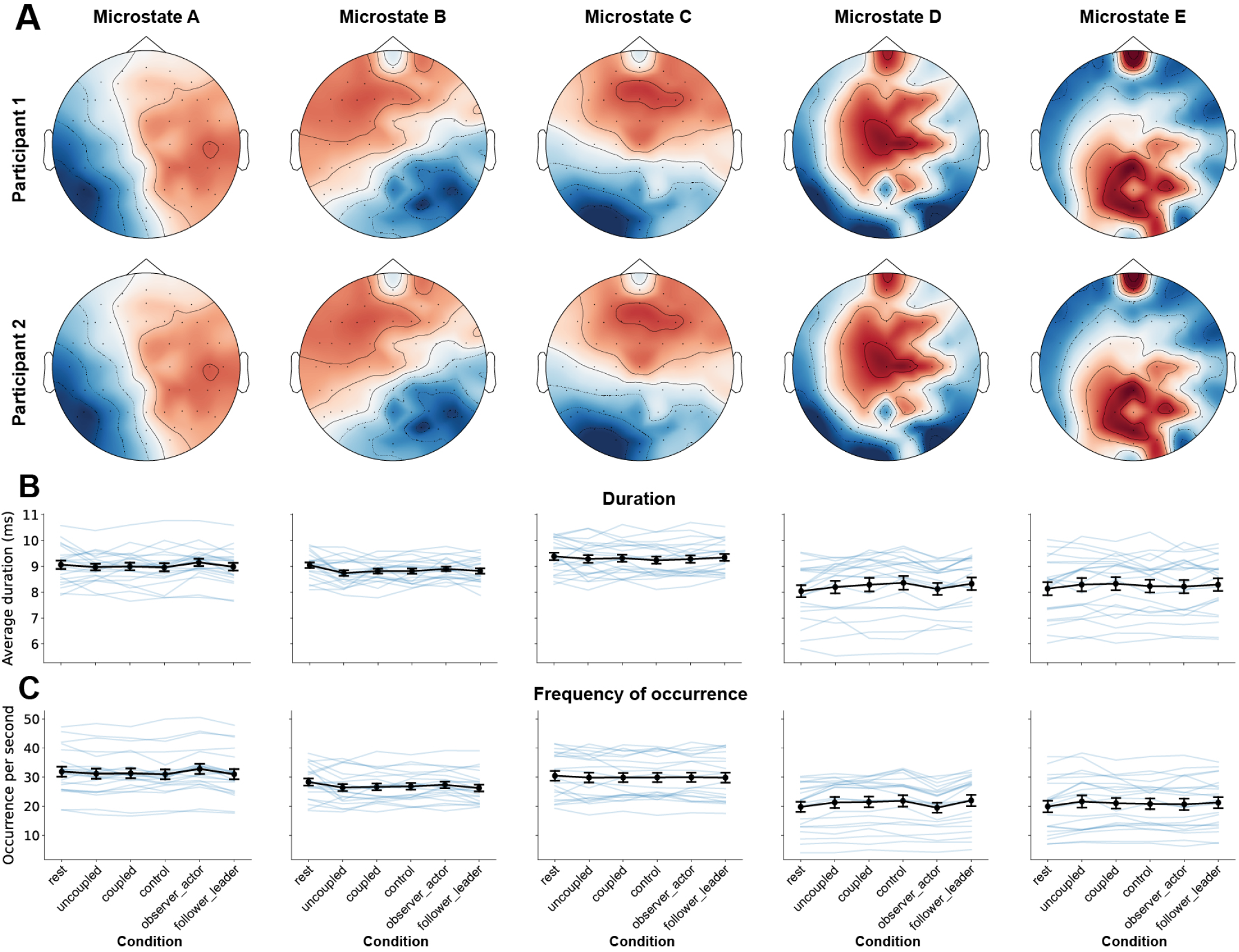
Inter-brain synchronization of alpha microstates. A) We investigated the co-occurrence of similar alpha microstates in the two participants in the dyads. B) The average duration the participants were in the same microstate. C) The frequency of co-occurrence of each microstate. None of the features were related to the different behavioural conditions.

**Fig. S11.**
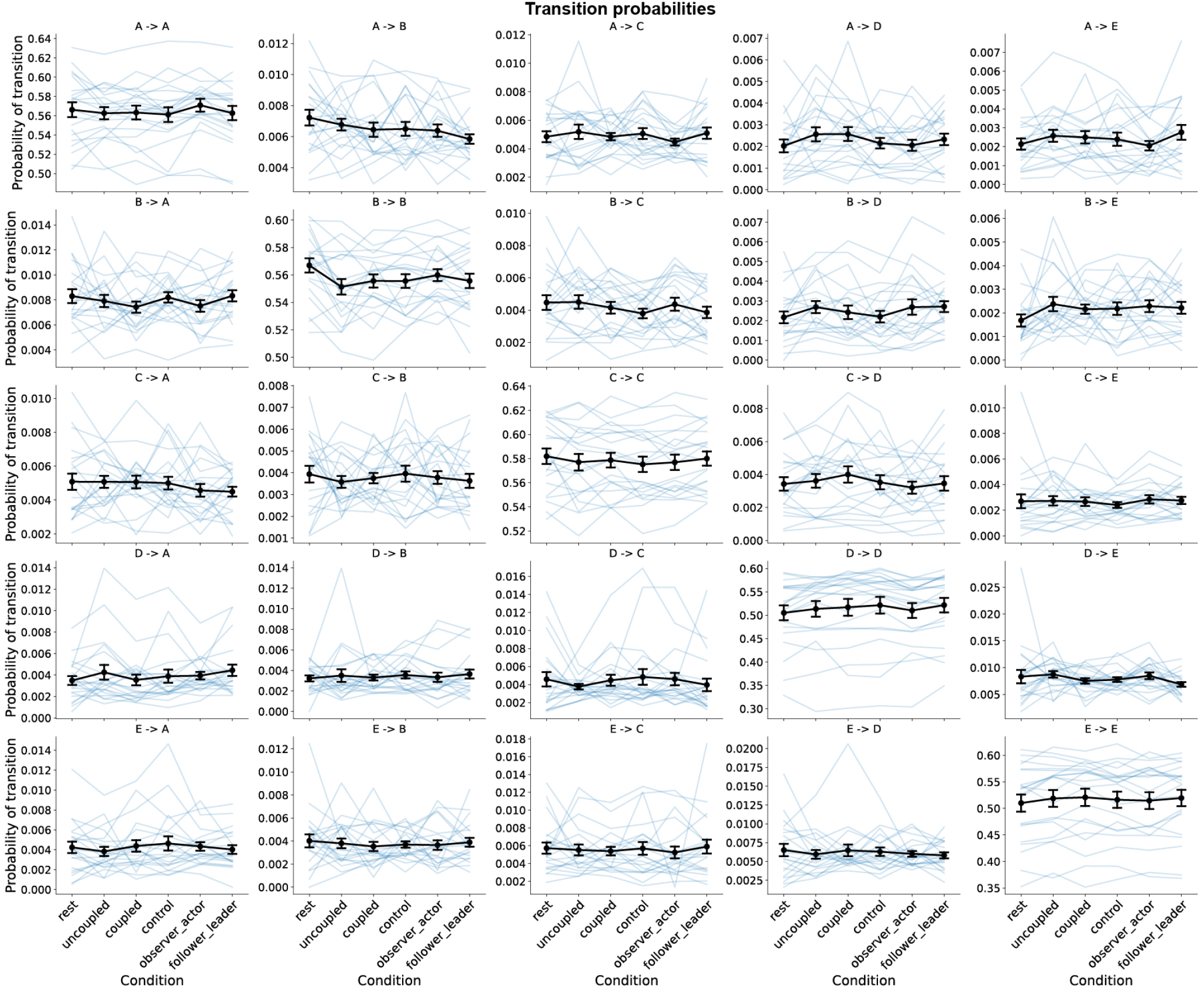
Inter-brain alpha microstate transition probabilities. The transition probabilities for co-occurring inter-brain alpha microstates. None of the features were related to the different behavioural conditions.

**Fig. S12.**
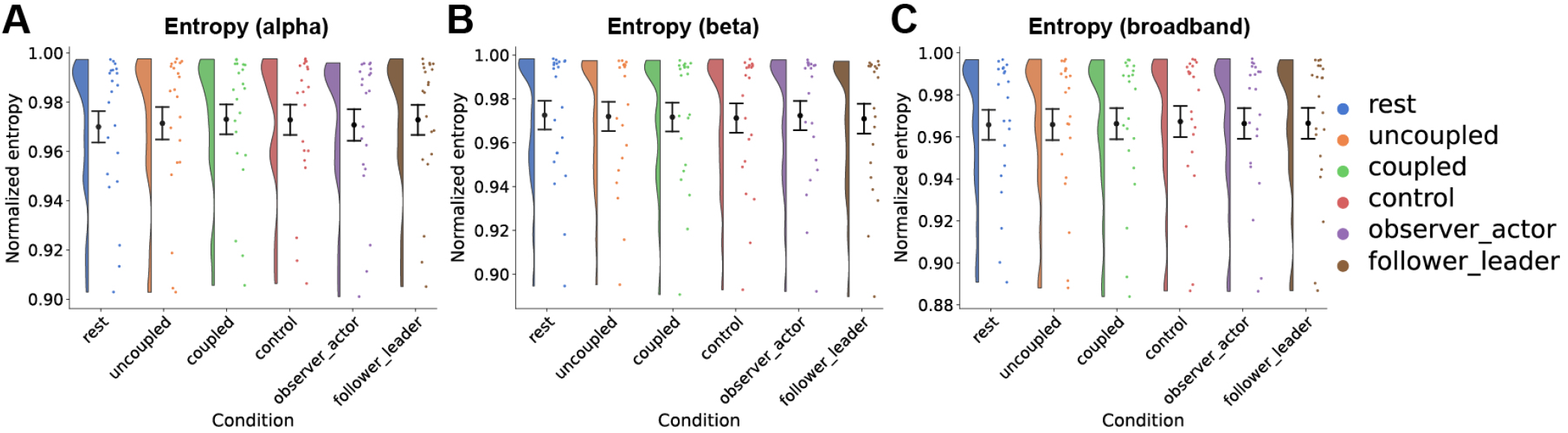
Inter-brain EEG microstate entropy. Shannon entropy was computed for the sequence time series for co-occurring microstates determined in the A) alpha, B) beta and C) broadband frequency range, normalized to the theoretical maximum (uniform label distribution). Entropy were not related to the different behavioural conditions.

**Fig. S13.**
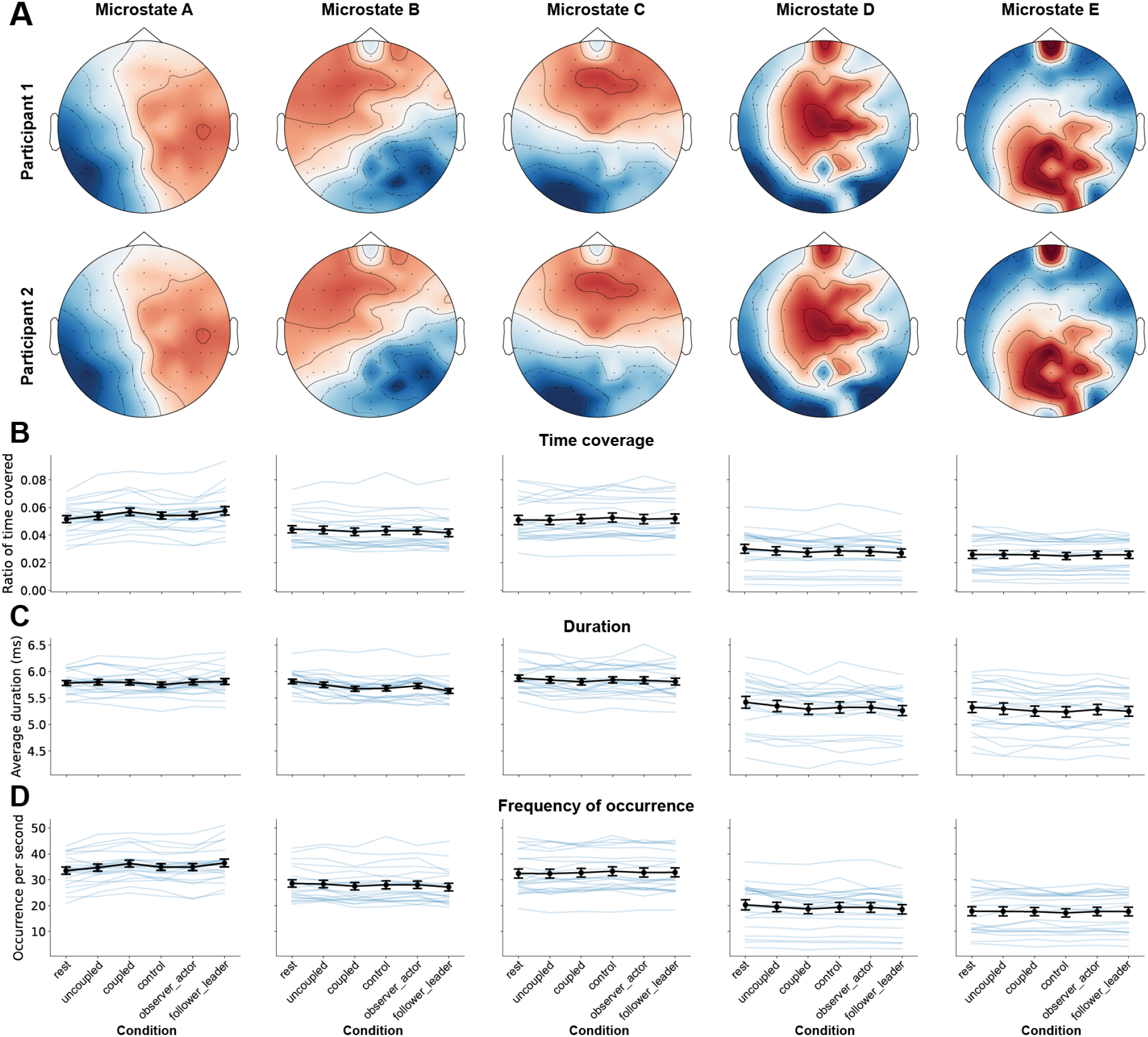
Inter-brain synchronization of beta microstates. A) We investigated the co-occurrence of similar beta microstates in the two participants in the dyads. B) The ratio of time the participants were in the same microstate. C) The average duration the participants were in the same microstate. D) The frequency of co-occurrence of each microstate. None of the features were related to the different behavioural conditions.

**Fig. S14.**
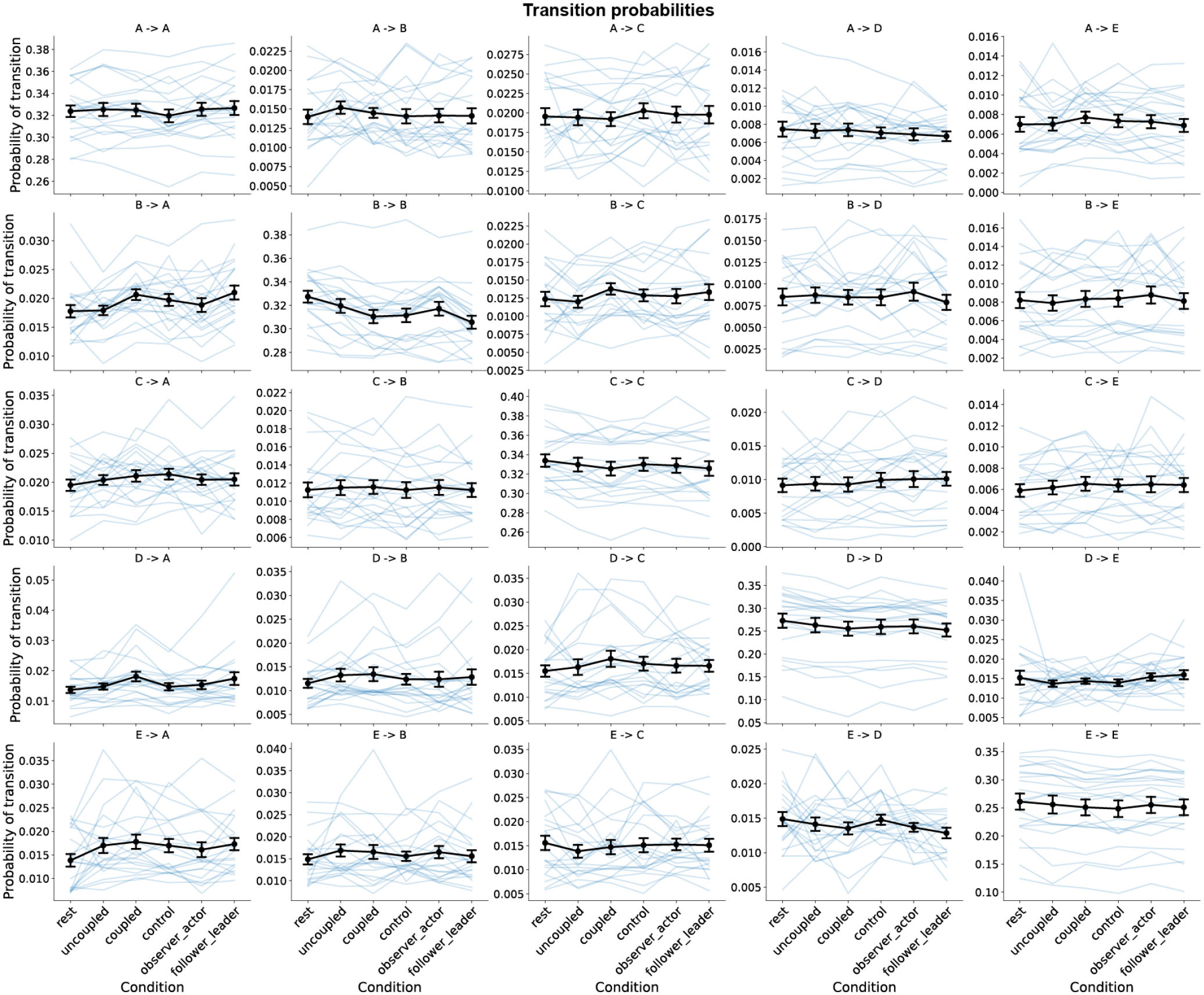
Inter-brain beta microstate transition probabilities. The transition probabilities for co-occurring inter-brain beta microstates. None of the features were related to the different behavioural conditions.

**Fig. S15.**
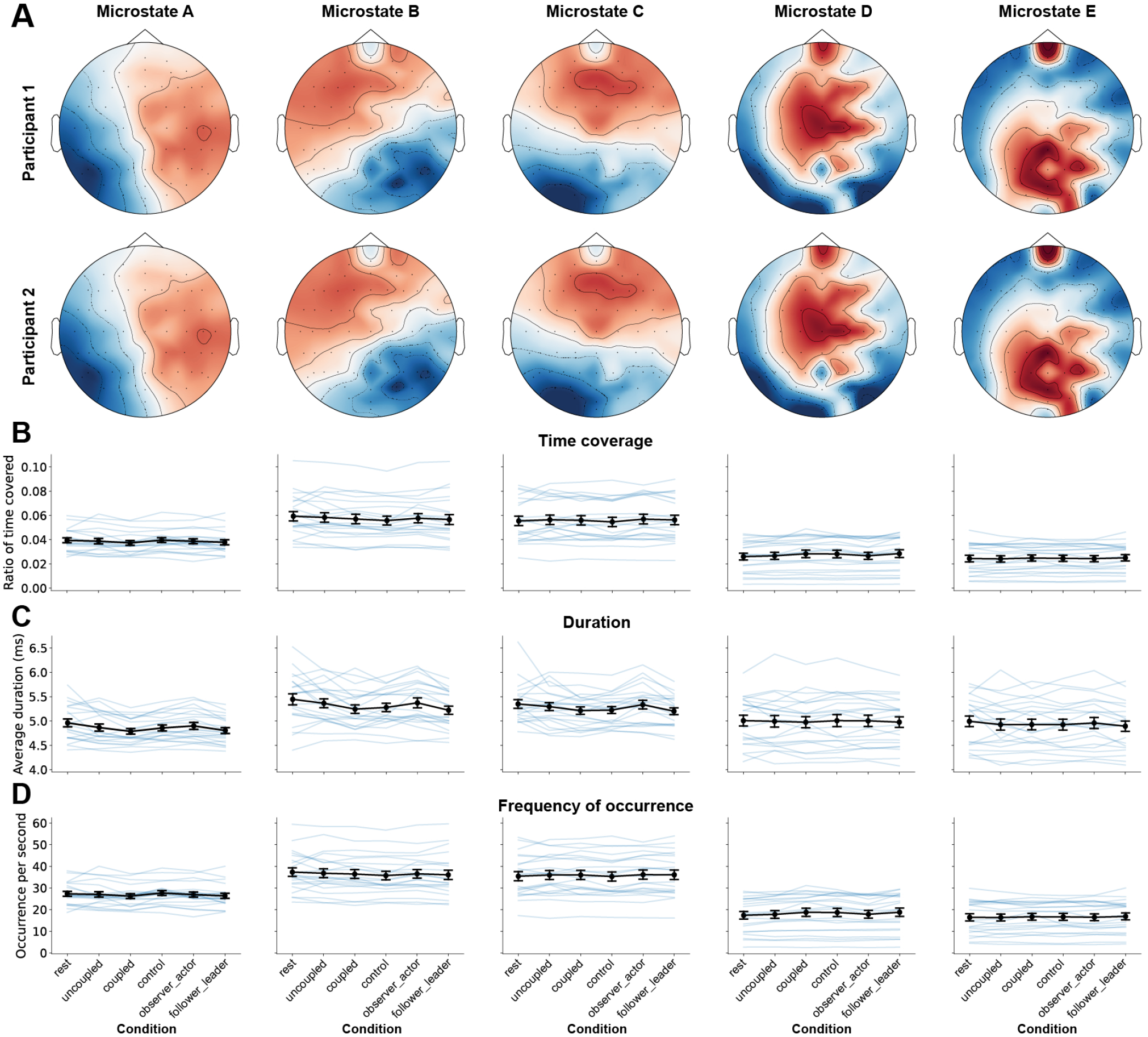
Inter-brain synchronization of broadband microstates. A) We investigated the co-occurrence of similar broadband microstates in the two participants in the dyads. B) The ratio of time the participants were in the same microstate. C) The average duration the participants were in the same microstate. D) The frequency of co-occurrence of each microstate. None of the features were related to the different behavioural conditions.

**Fig. S16.**
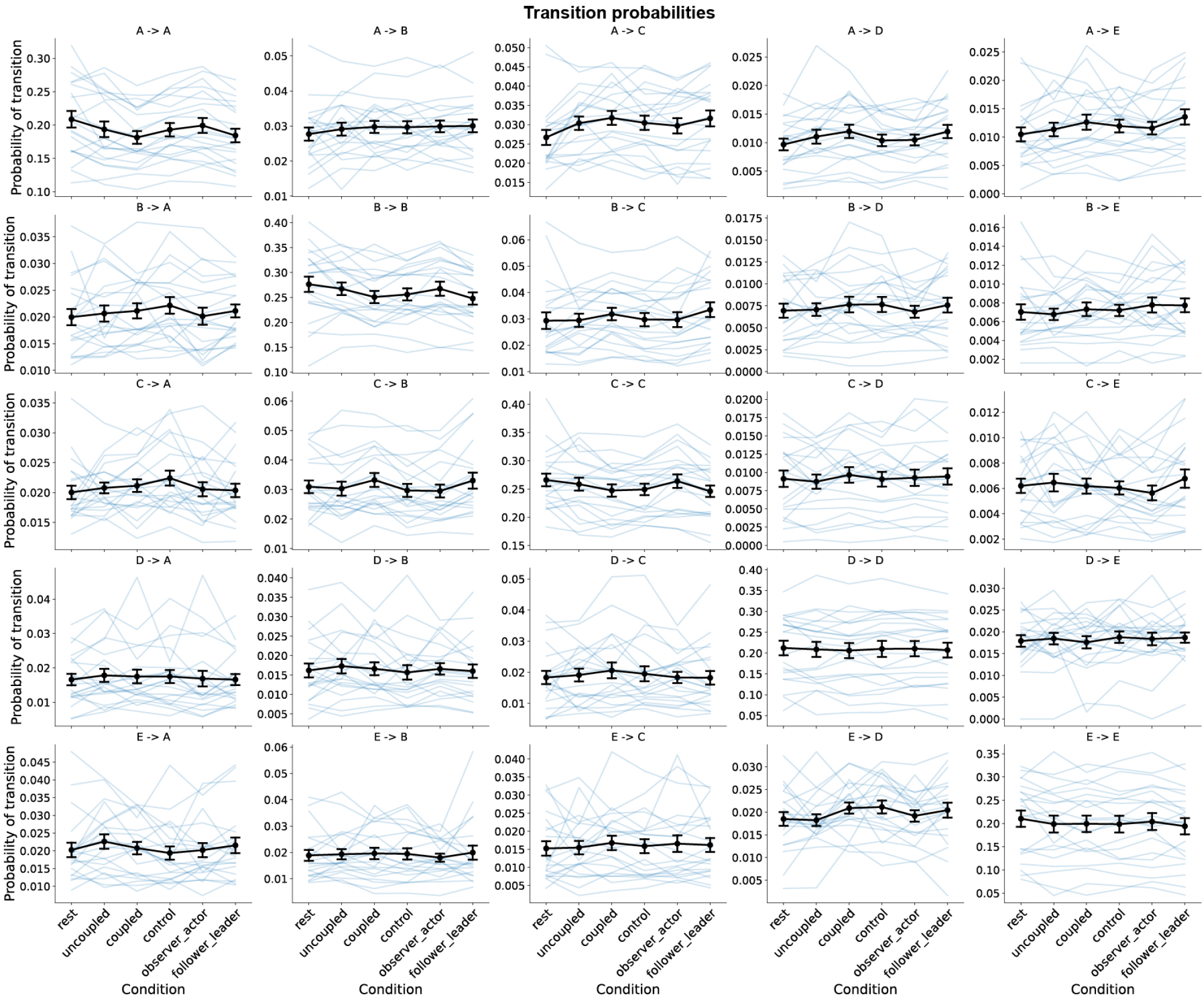
Inter-brain broadband microstate transition probabilities. The transition probabilities for co-occurring inter-brain broadband microstates. None of the features were related to the different behavioural conditions.

**Fig. S17.**
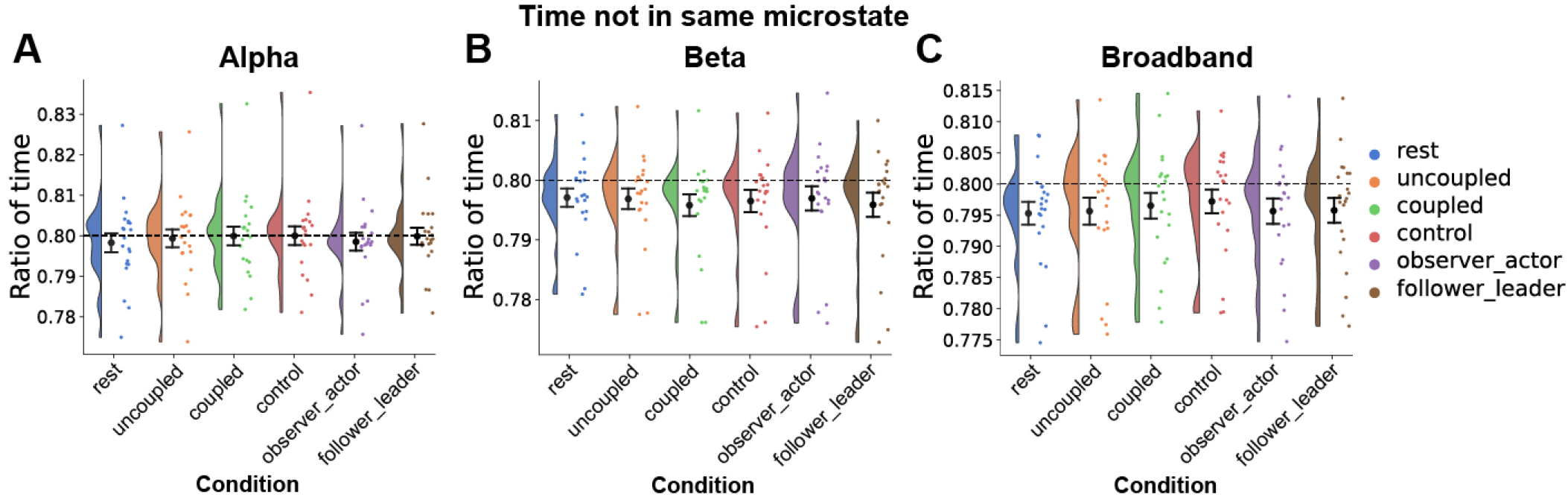
Ratio of time not in the same single-brain microstate. We explored the option of estimating interbrain synchronization as the time dynamics of when the two individuals in a pair were in the same single-brain microstate. However, the pairs were rarely both in the same microstate, in fact the amount of time they were in the same microstate was around chance-level for both A) alpha microstates, B) beta microstates and C) broadband microstates. The dashed line indicates 0.80, which correspond to chance-level with 5 microstates.

**Fig. S18.**
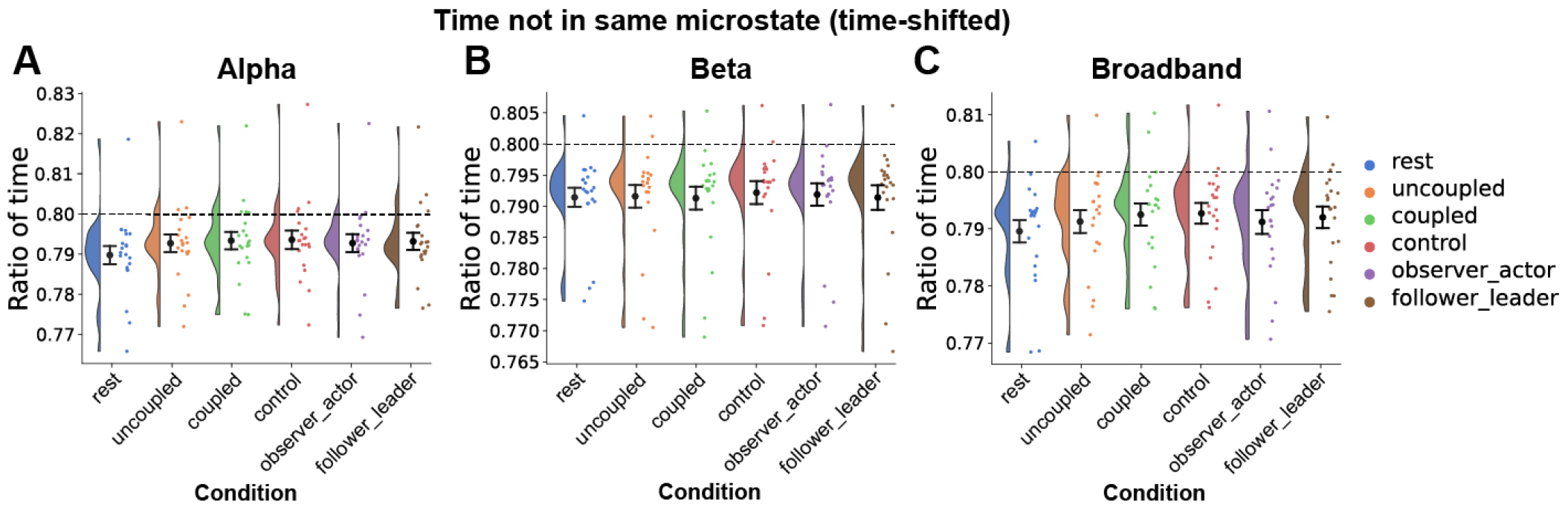
Ratio of time not in the same single-brain microstate, allowing for time-lags. We explored whether similar microstates at not precisely time-locked, but lagged timepoints, resulted in increased inter-brain synchronization. However, the decrease of ratio of time not in the same state was negligible for both A) alpha microstates, B) beta microstates and C) broadband microstates. The dashed line indicates 0.80, which correspond to chance-level with 5 microstates.

**Fig. S19.**
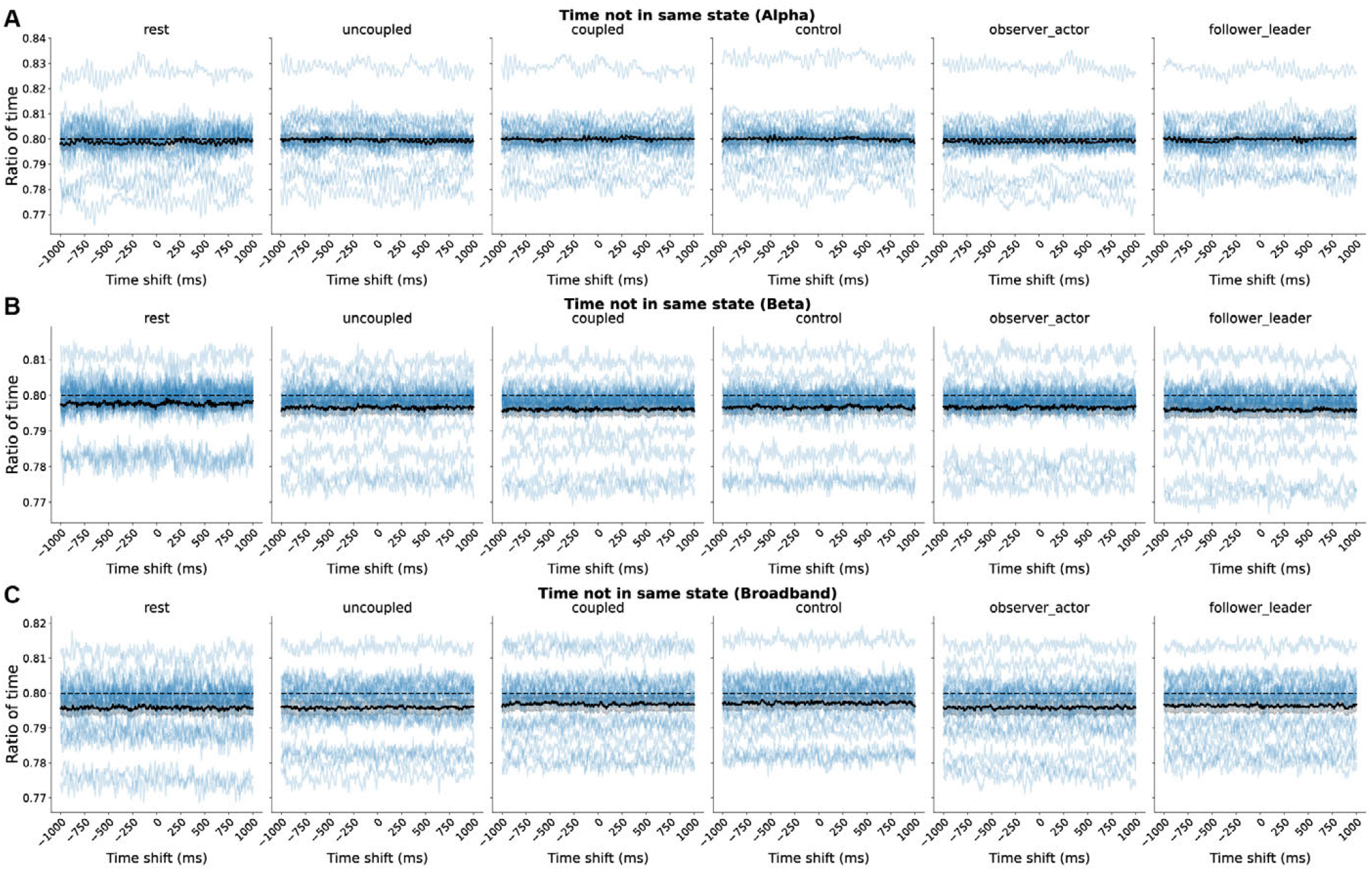
Ratio of time not in the same single-brain microstate at different time-lags. No clear time-lag resulted in an increase in time-lagged inter-brain synchronization (decrease in ratio of time not in the same microstate) for both A) alpha microstates, B) beta microstates and C) broadband microstates. The dashed line indicates 0.80, which correspond to chance-level with 5 microstates. The black line indicates the mean, with each blue line corresponding to each dyad.

**Fig. S20.**
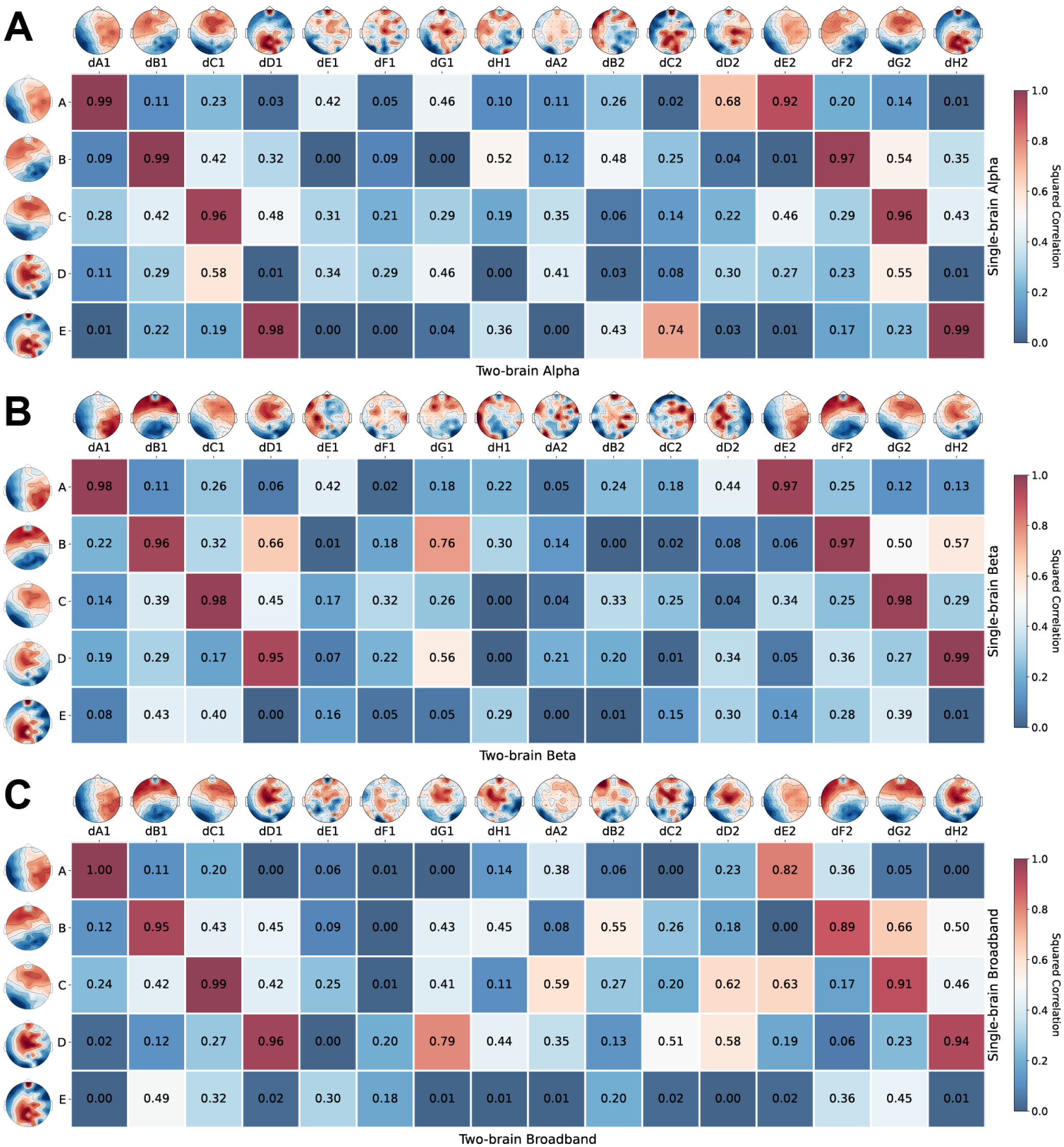
Spatial correlation between single-brain microstate and two-brain microstate topographies. The squared correlations between the single-brain and two-brain microstate topographies in A) alpha, B) beta, and C) broadband frequency ranges.

**Fig. S21.**
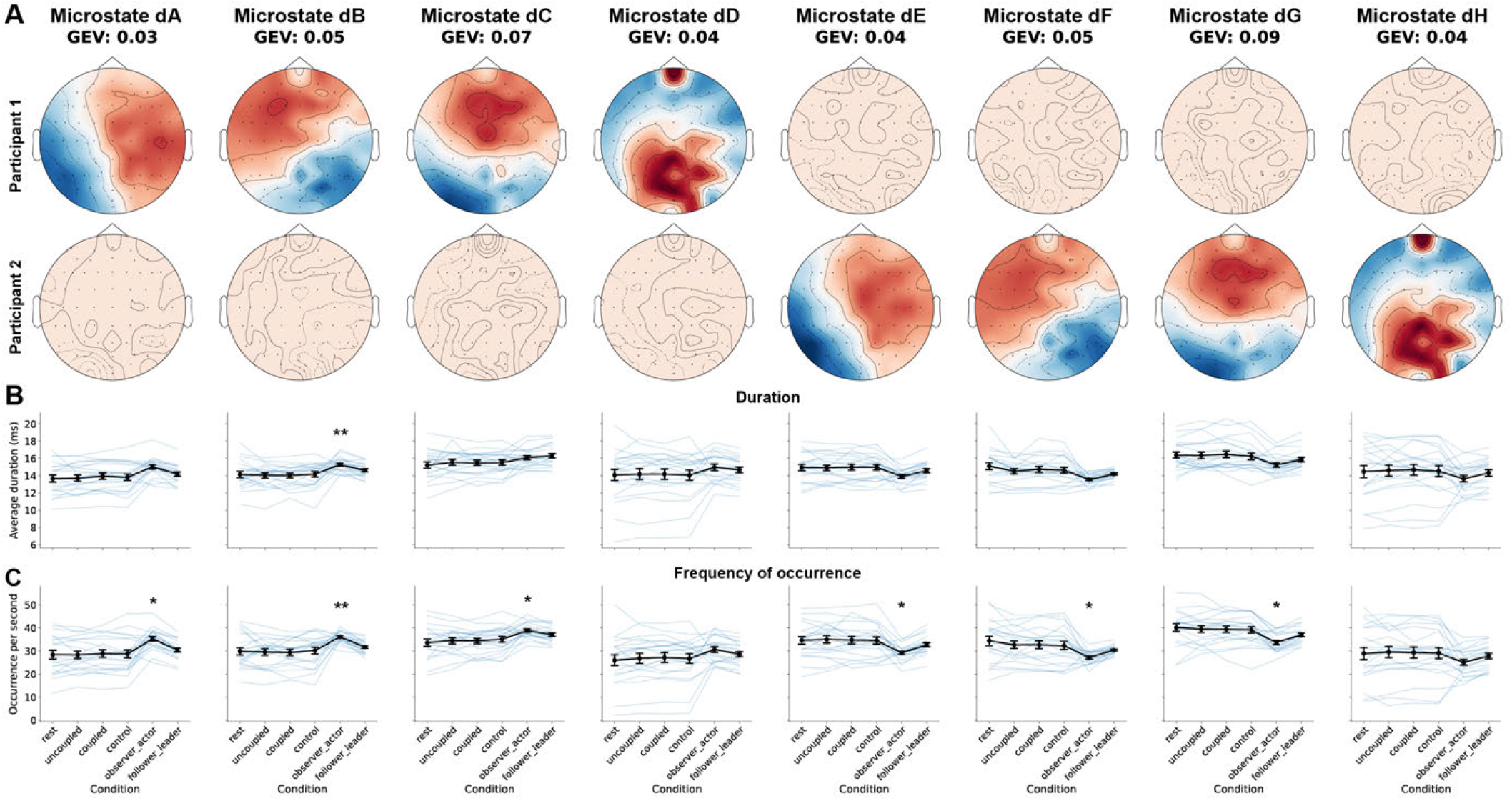
Two-brain alpha microstate analysis. A) When the two-brain alpha microstates were plotted using a fixed color scale based on the min and max value across all the microstates, it became evident that dA2, dB2, dC2, dD2, dE1, dF1, dG1 and dH1 reflected the mean topography, with a z-scored activity around 0. B) The average duration of each two-brain microstate. C) The frequency of occurrence for each two-brain microstate. Two-brain microstate dynamics were different between the asymmetrical observer-actor and symmetrical conditions. GEV: global explained variance.

**Fig. S22.**
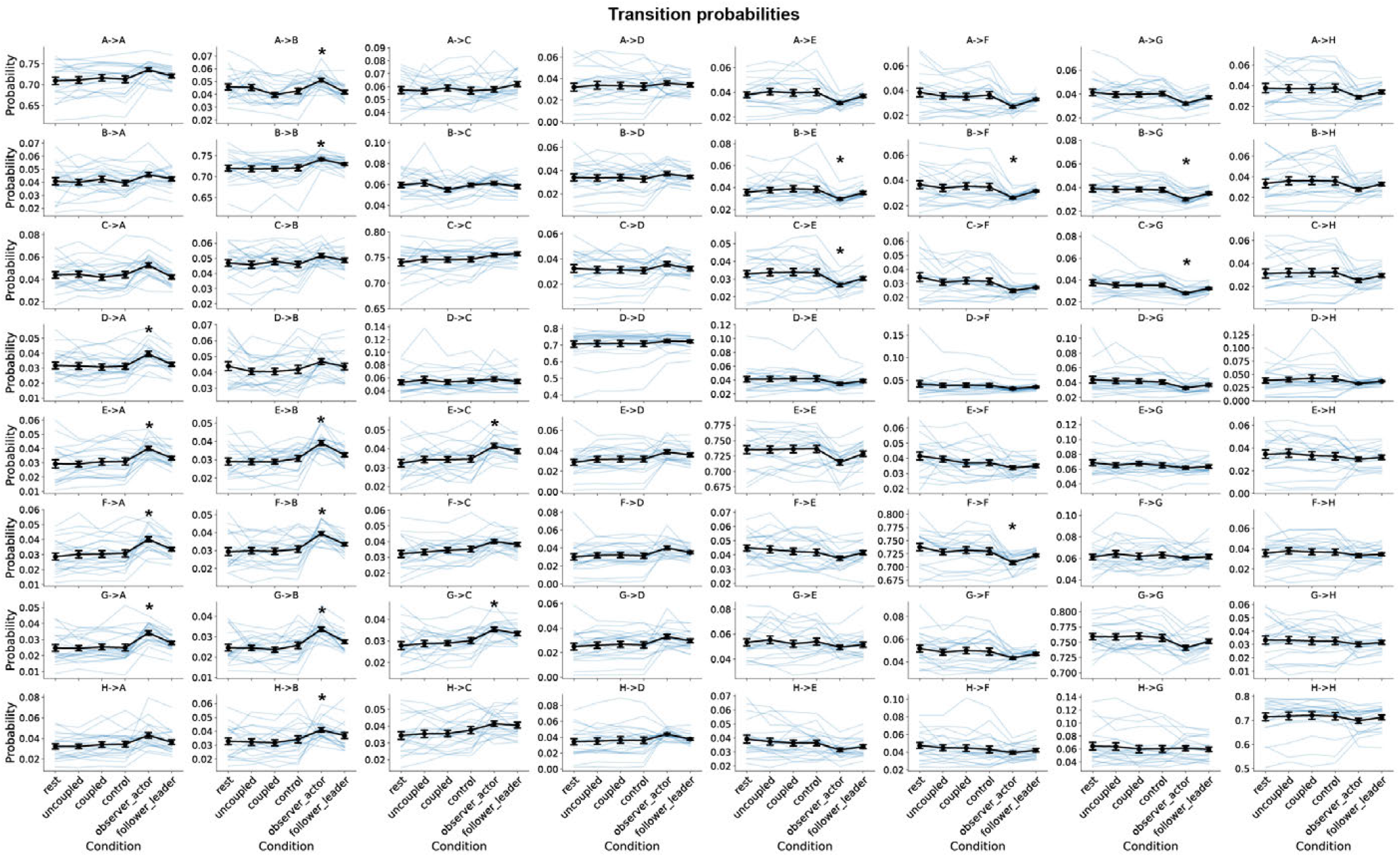
Two-brain alpha microstate transition probabilities. Two-brain alpha microstate transition probabilities were different between the asymmetrical and symmetrical conditions, with only the observer-actor condition differing after multiple-comparison correction.

**Fig. S23.**
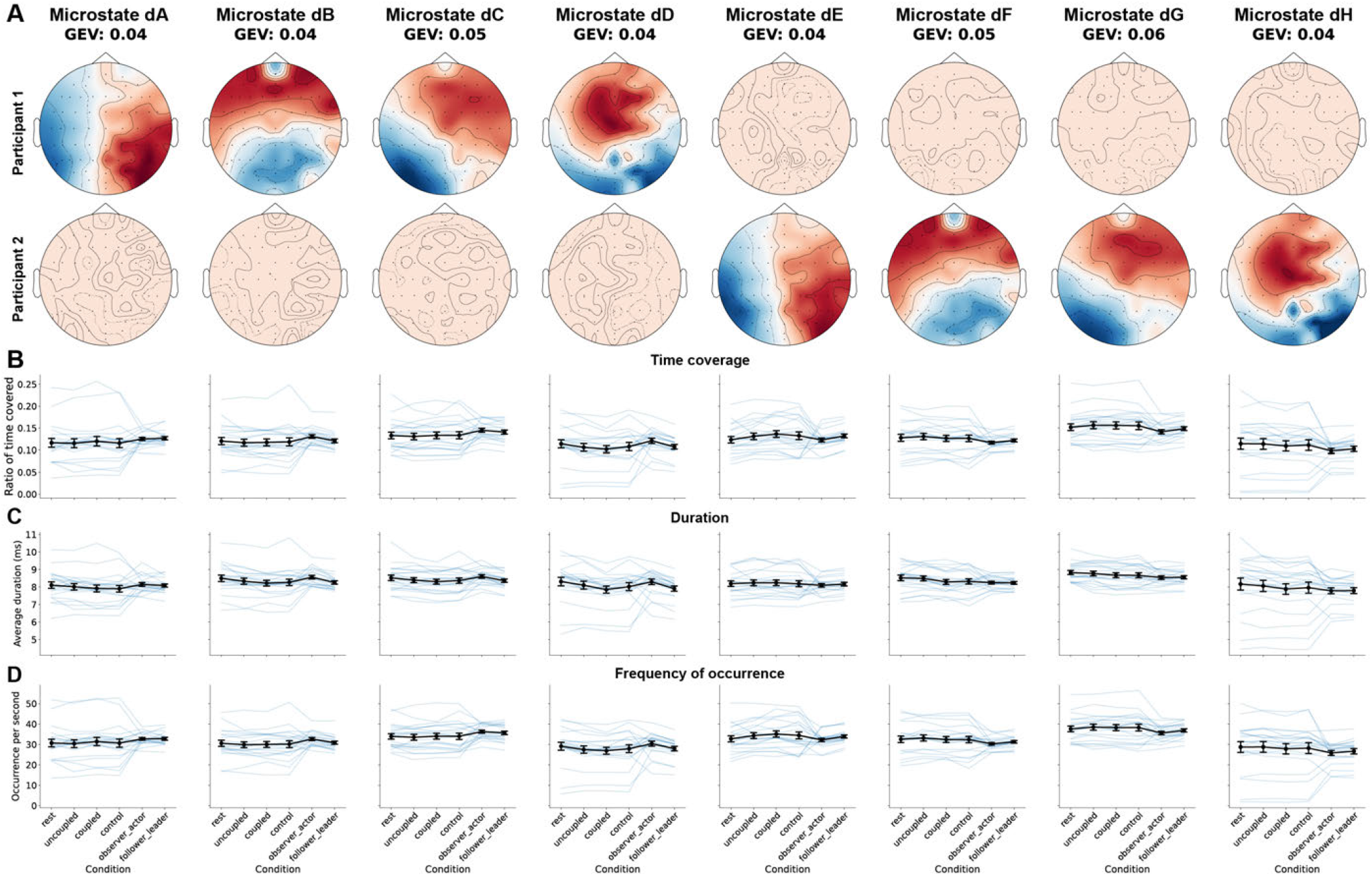
Two-brain beta microstate analysis. A) Microstate analysis was performed on the simultaneously recorded EEG from 21 pairs collected during the mirror-game paradigm, which yielded eight two-brain beta microstates, explaining around 36% of the variance. B) The ratio of time covered by each two-brain microstate. C) The average duration of each two-brain microstate. D) The frequency of occurrence for each two-brain microstate. None of the features were related to the different behavioural conditions. GEV: global explained variance.

**Fig. S24.**
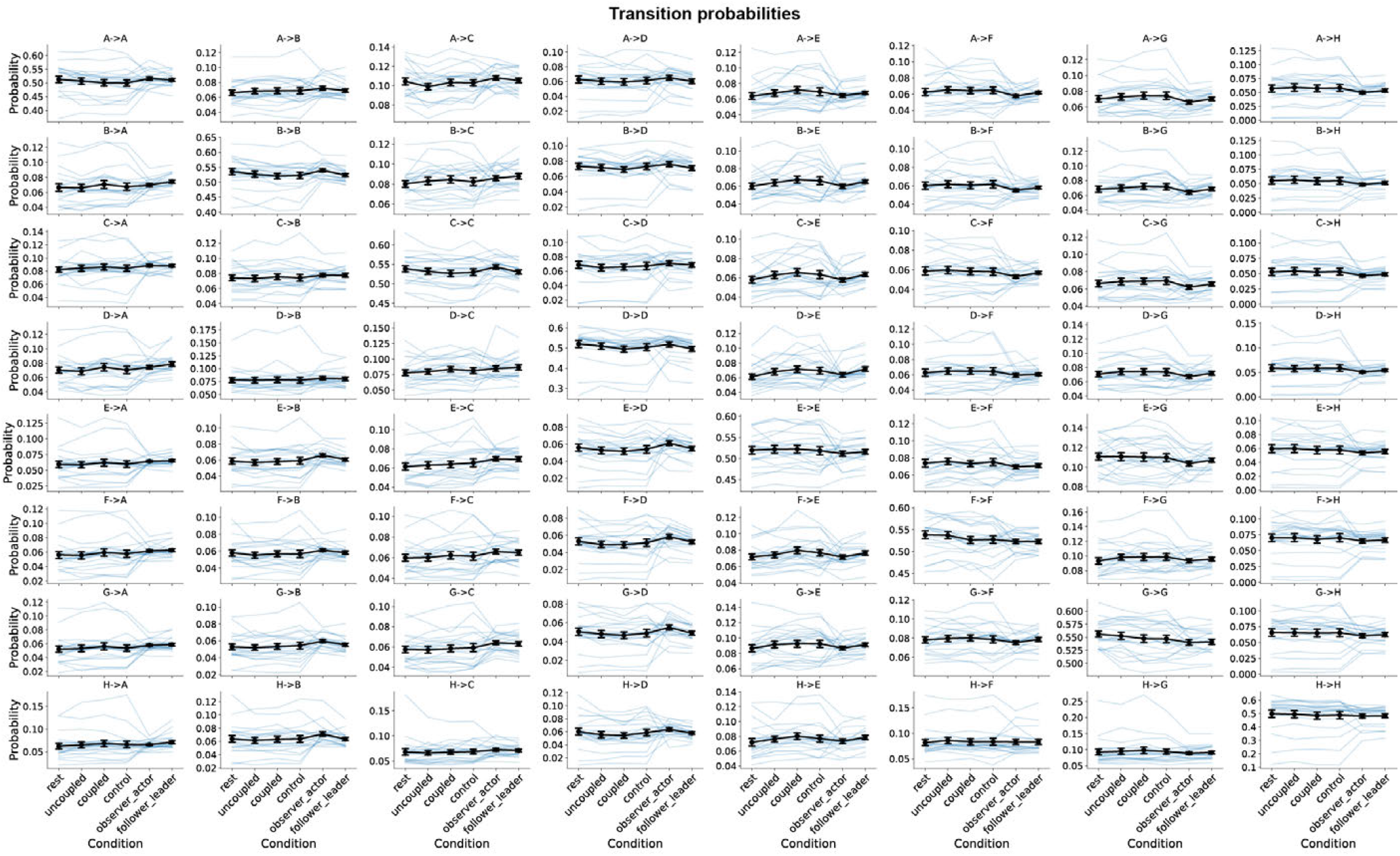
Two-brain beta microstate transition probabilities. The probabilities of transitioning between each two-brain beta microstate were not related to the different behavioural conditions after multiple-comparison correction.

**Fig. S25.**
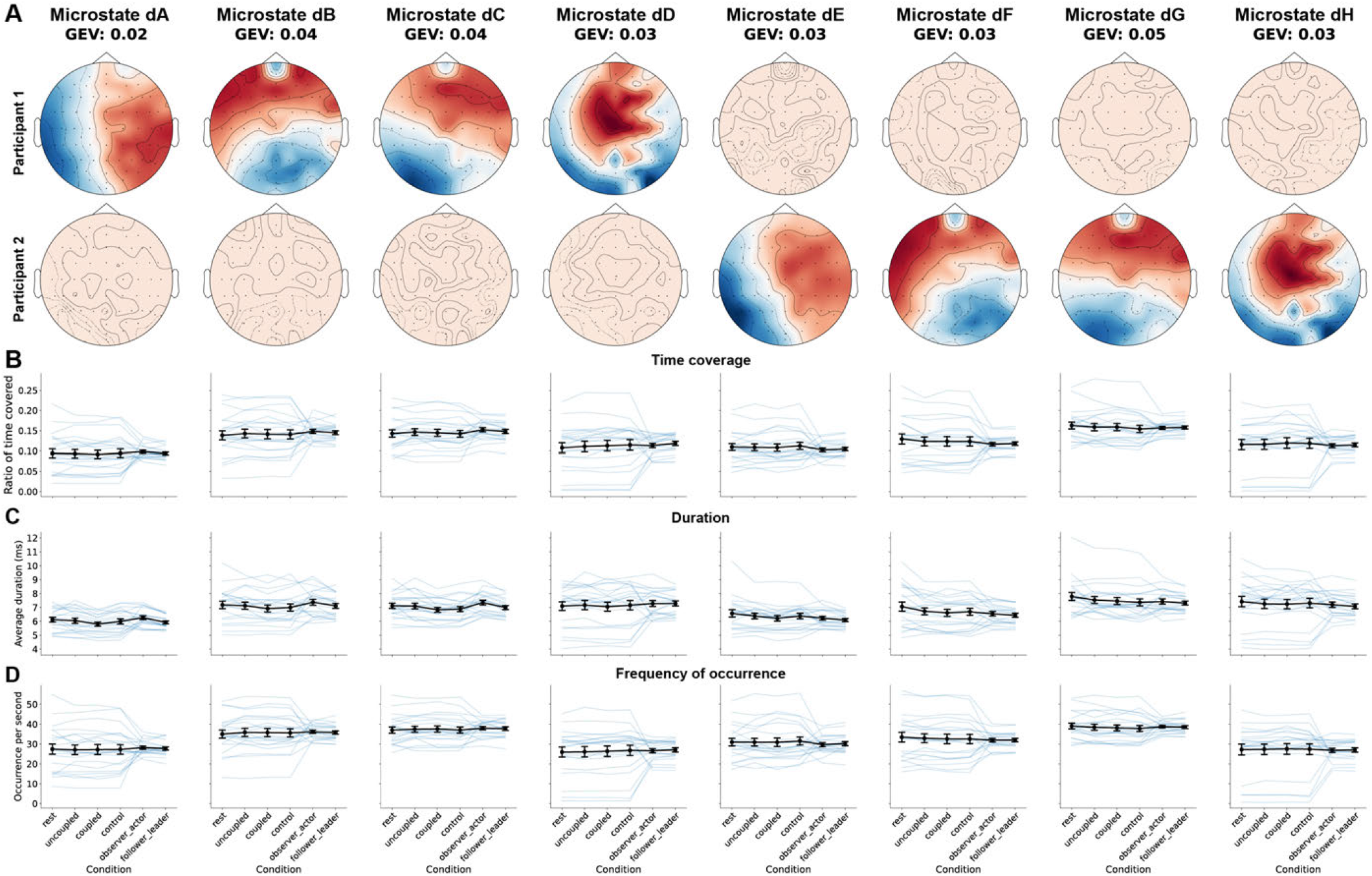
Two-brain broadband microstate analysis. A) Microstate analysis was performed on the simultaneously recorded EEG from 21 pairs collected during the mirror-game paradigm, which yielded eight two-brain broadband microstates, explaining around 27% of the variance. B) The ratio of time covered by each two-brain microstate. C) The average duration of each two-brain microstate. D) The frequency of occurrence for each two-brain microstate. None of the features were related to the different behavioural conditions after multiple-comparison correction. GEV: global explained variance.

**Fig. S26.**
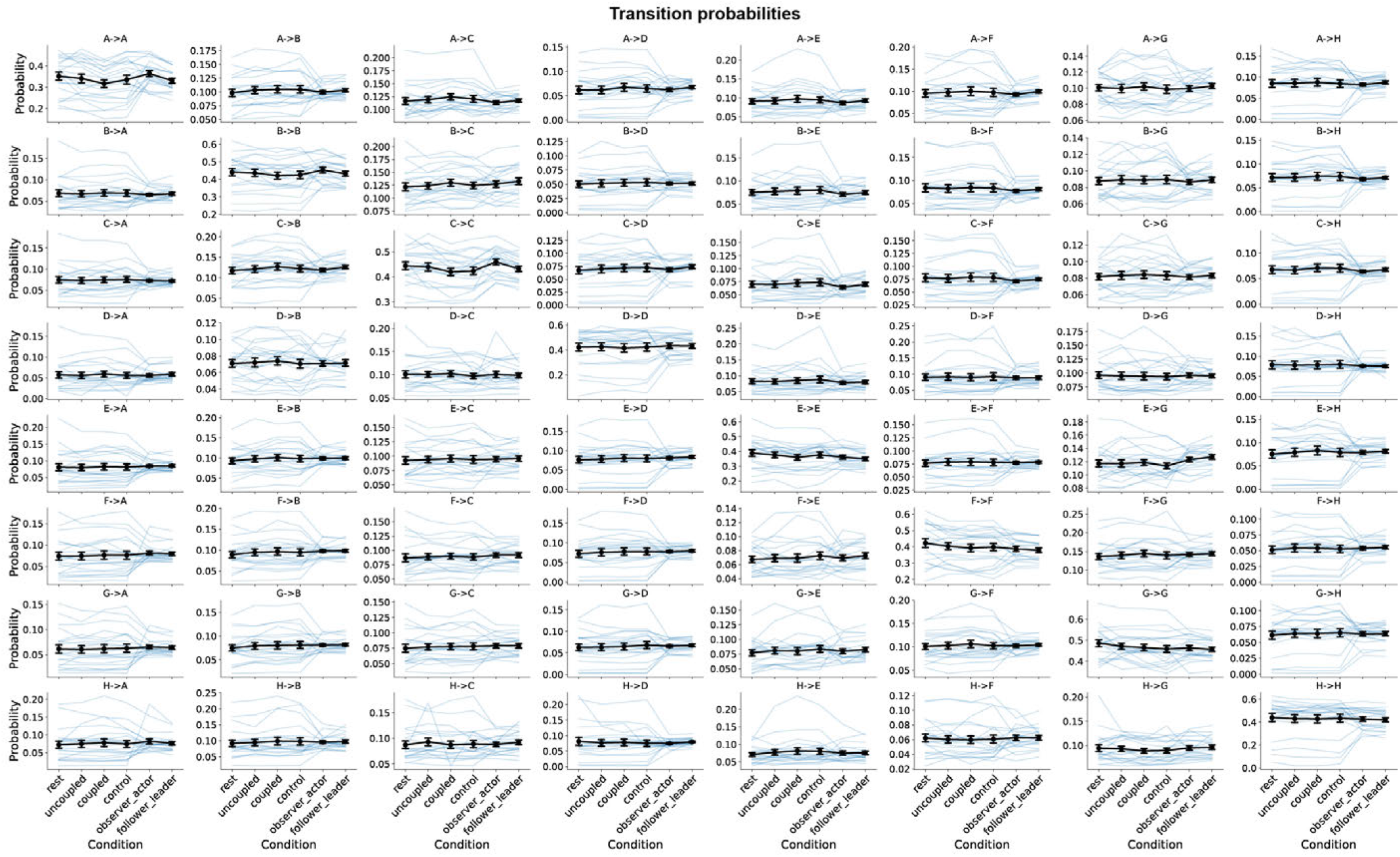
Two-brain broadband microstate transition probabilities. The probabilities of transitioning between each two-brain broadband microstate were not related to the different behavioural conditions after multiple-comparison correction.

**Fig. S27.**
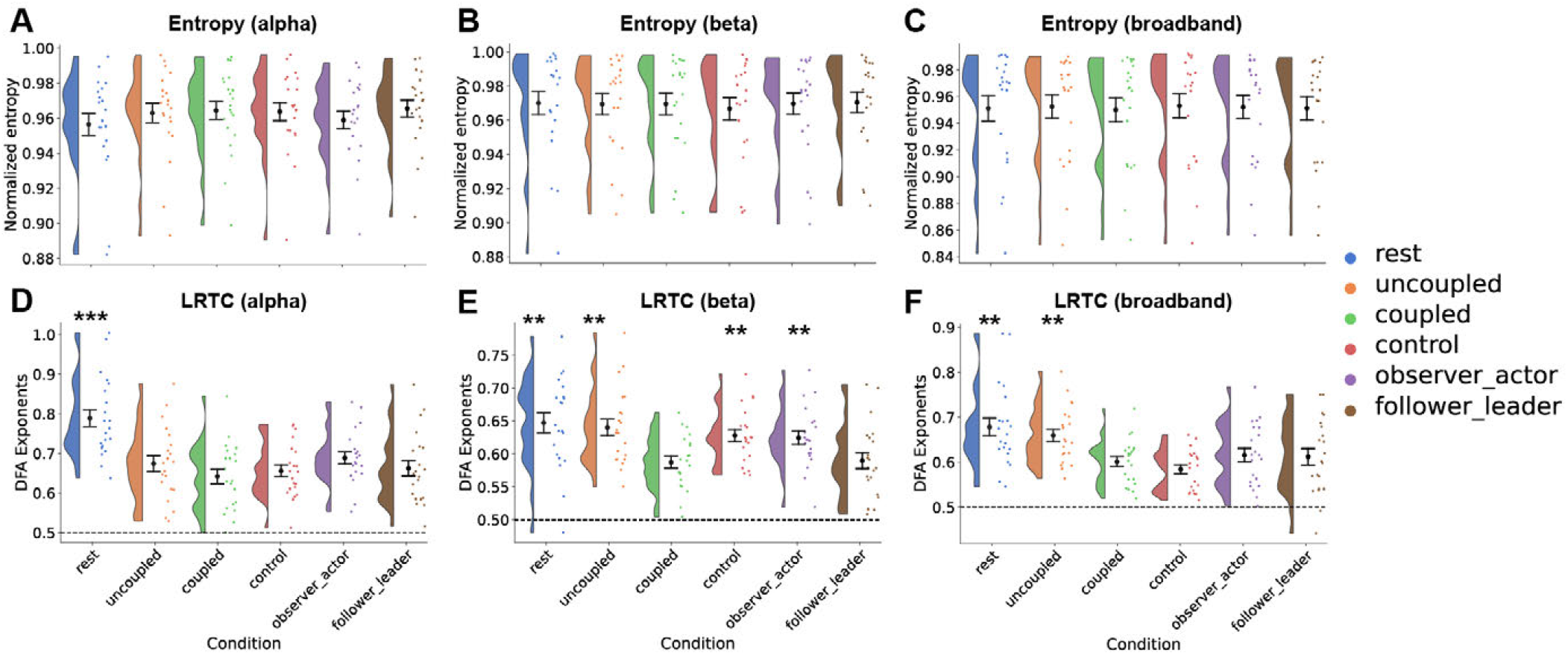
Complexity measures for two-brain EEG microstates. Shannon entropy was computed for the microstate sequence time series, normalized to the theoretical maximum (uniform label distribution), for the A) alpha, B) beta and C) broadband frequency range. Long-range temporal correlations in the form of Hurst exponents were estimated as DFA exponents in the D) alpha, E) beta and F) broadband frequency range. Entropy features were not related to the different behavioural conditions, however, the rest condition had significantly higher DFA exponents than the baseline coupled condition for all three frequency ranges, while uncoupled also had higher DFA exponents than coupled in the beta and broadband frequency range. The control and observer-actor condition was also associated with higher DFA exponents than the coupled condition in the beta frequency range. To perform DFA on the microstate sequence time series, dA, dB, dC and dD was partitioned into class 1 and dE, dF, dG and dH into class 2. GEV: global explained variance, DFA: detrended fluctuation analysis, LRTC: long-range temporal correlations.

**Fig. S28.**
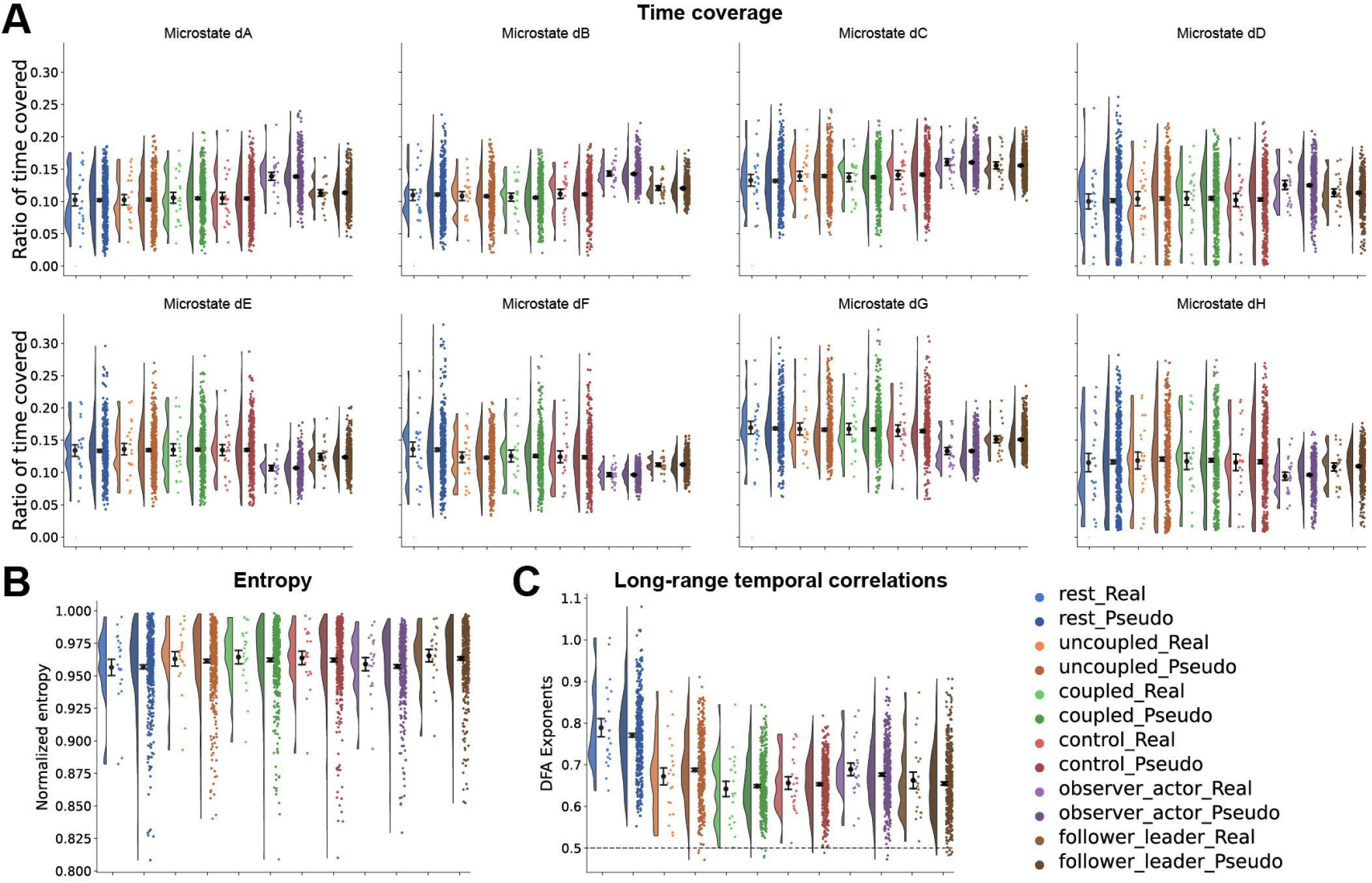
Two-brain microstate dynamics were similar between real pairs and pseudo-pairs. The features shown are for the alpha frequency band. A) The time coverage was greater in dA, dB, dC and dD and lower in dE, dF, dG and dH in the asymmetrical tasks compared to principal symmetric coupled condition, however the difference was similar between the real pairs and the pseudo-pairs. The microstate sequence time series of the pseudo-pairs also had similar B) entropy and C) longrange temporal correlation as the real pairs. The colors correspond to each condition, with darker shades corresponding to the pseudo-pairs.

**Fig. S29.**
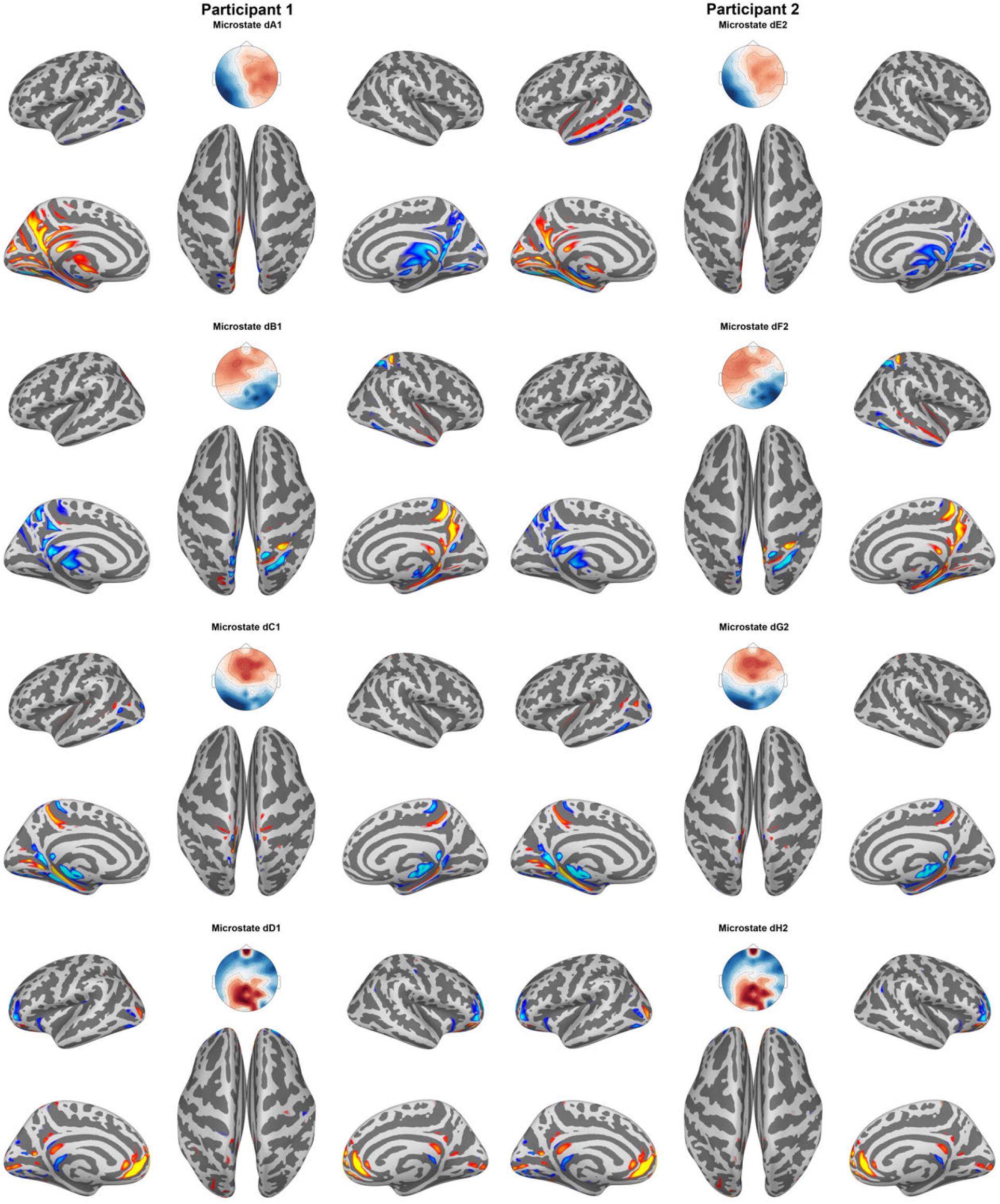
The eight two-brain alpha microstates and their corresponding eLORETA cortical activities. We only showed the cortical sources for dA1, dB1, dC1, dD1, dE2, dF2, dG2 and dH2 as the other two-brain microstates reflected the mean arbitrary activity (i.e. had no noticeable areas of activity).

**Fig. S30.**
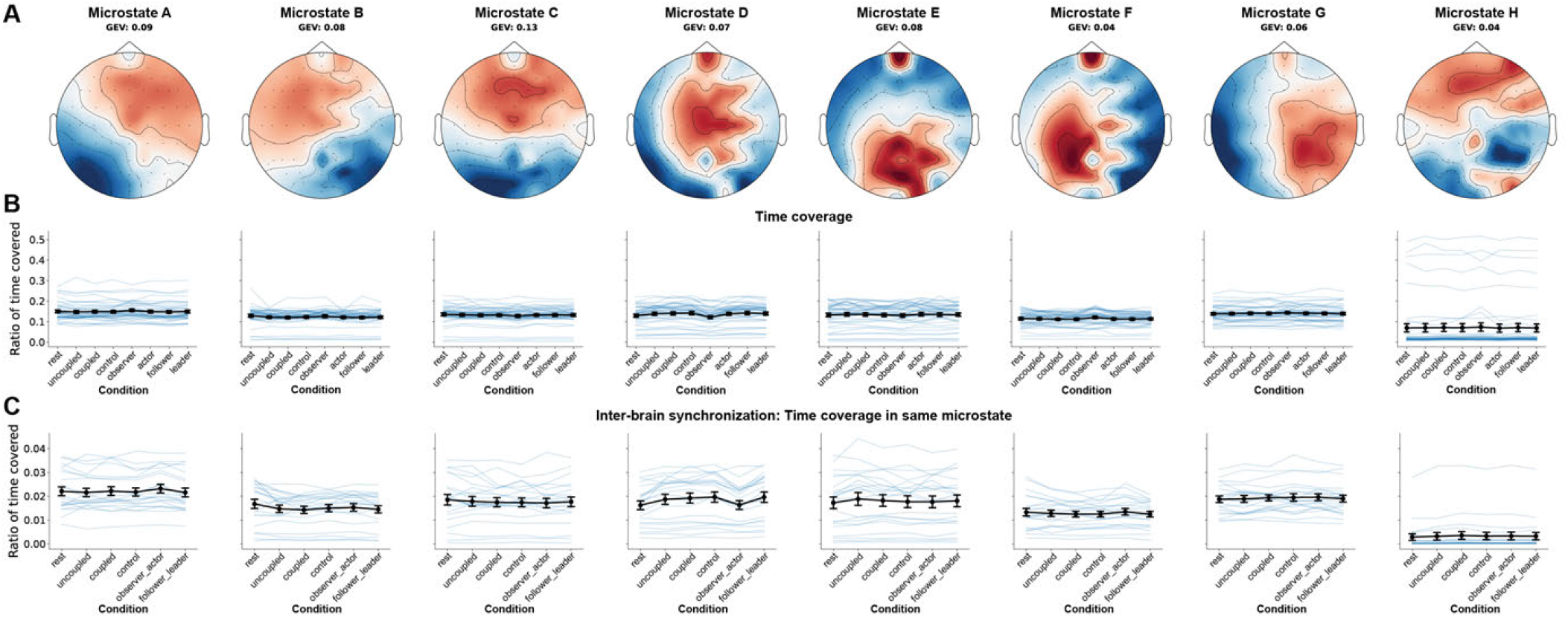
Single-brain EEG microstate analysis with eight extracted microstates. A) To be consistent with the eight extracted microstates for the two-brain microstate analysis, we repeated the single-brain microstate analysis using the same cluster number. In total, the eight microstates explained around 60% of the variance. B) The ratio of time covered by each microstate was not related to the different behavioural conditions. C) The ratio of time the two participants in a dyad were in the same microstate was also not related to the different behavioural conditions. Notice the ratio of time coverage in C) was normalized to the total time they were in the same microstate and the time not in the same microstate (Supplementary Figure S17). GEV: global explained variance.

**Fig. S31.**
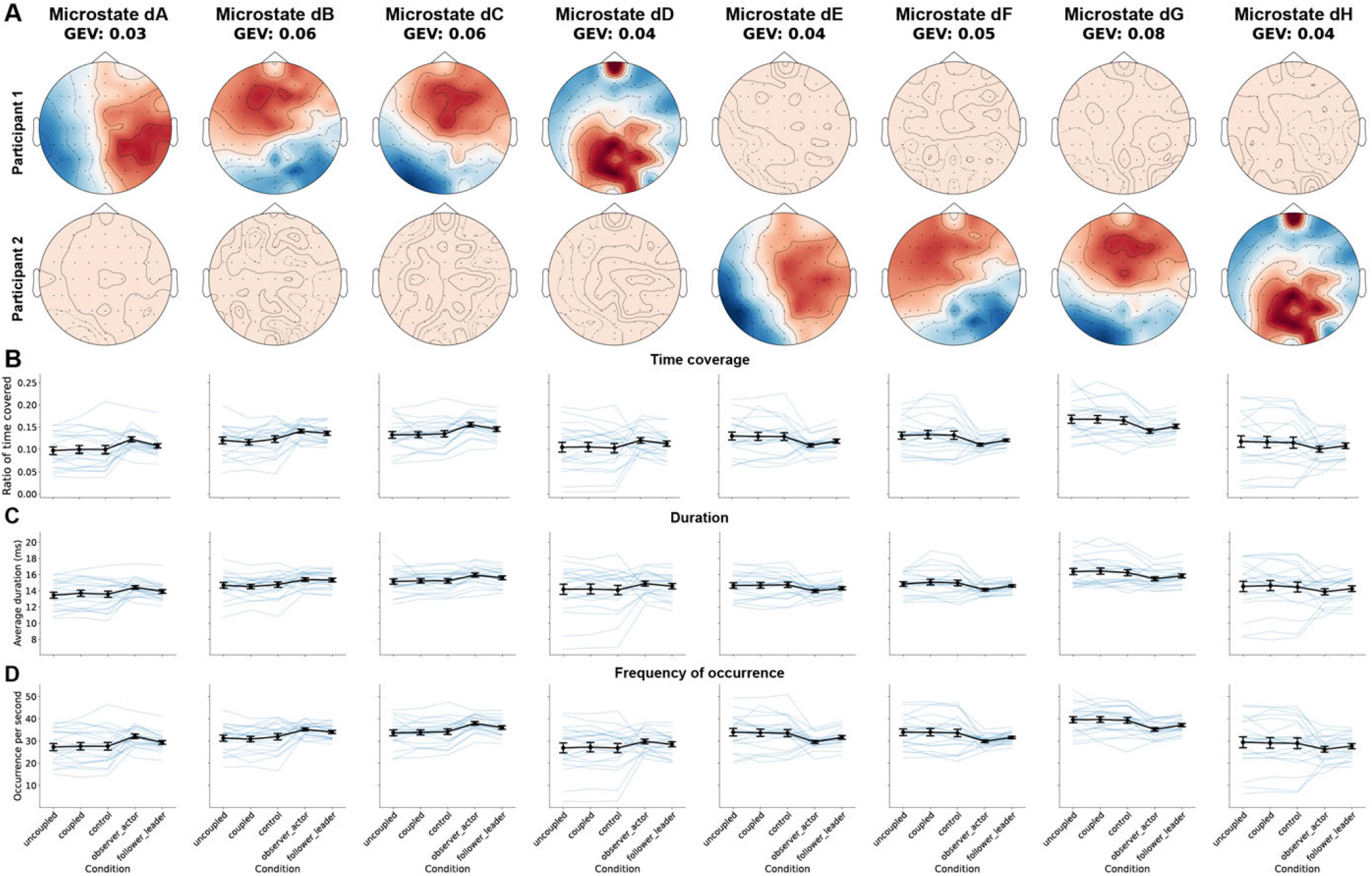
Removing rest conditions did not change the determined two-brain microstates. A) We repeated the two-brain alpha microstate fitting after excluding the rest conditions. No clear qualitative differences were observed. B) The ratio of time covered by each two-brain microstate fitted without rest conditions showed the same trends with greater time coverage of canonical resting-state-like microstates in the observer (dA, dB, dC and dD) and lower time coverage in the observer (dE, dF, dG and dH). GEV: global explained variance.

